# The Radiogenomic and Spatiogenomic Landscapes of Glioblastoma, and their Relationship to Oncogenic Drivers

**DOI:** 10.1101/2022.12.15.517767

**Authors:** Anahita Fathi Kazerooni, Hamed Akbari, Xiaoju Hu, Vikas Bommineni, Dimitris Grigoriadis, Erik Toorens, Chiharu Sako, Elizabeth Mamourian, Dominique Ballinger, Robyn Sussman, Ashish Singh, Ioannis I. Verginadis, Nadia Dahmane, Constantinos Koumenis, Zev A. Binder, Stephen J. Bagley, Suyash Mohan, Artemis Hatzigeorgiou, Donald M. O’Rourke, Tapan Ganguly, Subhajyoti De, Spyridon Bakas, MacLean P. Nasrallah, Christos Davatzikos

**Author notes:** Equal contribution (MacLean P. Nasrallah, Christos Davatzikos). Correspondence. Christos Davatzikos, PhD Address: 3700 Hamilton Walk, 7th floor, Philadelphia PA 191045. **Author contributions.** *Conception and design: A Fathi Kazerooni, H Akbari, X Hu, S De, D Grigoriadis, A Hatzigeorgiou, S Bakas, M Nasrallah, C Davatzikos* Development of methodology: A Fathi Kazerooni, H Akbari, X Hu, D Grigoriadis, V Bommineni, E Toorens, T Ganguly, S Bakas, M Nasrallah, C Davatzikos Acquisition of data: C Sako, E Mamourian, E Toorens, T Ganguly, R Sussman, S Bakas Preprocessing of data: C Sako, E Mamourian, A Fathi Kazerooni, E Toorens Analysis and interpretation of data (e.g., statistical analysis, biostatistics, computational analysis): A Fathi Kazerooni, H Akbari, X Hu, S De, D Grigoriadis, M Nasrallah, C Davatzikos Writing, review, and/or revision of the manuscript: A Fathi Kazerooni, H Akbari, X Hu, V Bommineni, D Grigoriadis, E Toorens, S Bagley, C Sako, E Mamourian, D Ballinger, R Sussman, A Singh, I Verginadis, N Dahmane, C Koumenis, Z A Binder, S J Bagley, S Mohan, A Hatzigeorgiou, D M. O’Rourke, T Ganguly, S De, S Bakas, M Nasrallah, C Davatzikos.

## Abstract

Glioblastoma (GBM) is well-known for its molecular and spatial heterogeneity, which poses a challenge for precision therapies and clinical trial stratification. Here, in a comprehensive radiogenomics study of 358 GBMs, we investigated the associations between the imaging and spatial characteristics of the tumors with their cancer gene mutation status, as well as with the cross-sectionally inferred likely order of mutational events. We show that cross-validated machine learning analysis of multi-parametric MRI scans results in distinctive *in vivo* imaging signatures of several mutations, which are relatively more distinctive in homogeneous tumors which harbor only one of these mutations. These imaging signatures offer mechanistic insights into how various mutations influence the phenotype of the tumor and its surrounding infiltrated brain tissue via neovascularization and vascular leakage, increased cell density, invasion and migration, and other characteristics captured by respective imaging features. Furthermore, we found that spatial location and tumor distribution vary, depending on the GBM’s molecular characteristics. Finally, distinct imaging and spatial characteristics were associated with cross-sectionally estimated evolutionary trajectories of the tumors. Collectively, our study establishes a panel of *in vivo* and clinically accessible imaging-AI biomarkers of GBM that reflect their molecular composition and oncogenic drivers.

## 1 Introduction

Most cancers harbor a constellation of tumor subpopulations with highly variable genetic and epigenetic properties, giving rise to spatial and molecular intratumoral heterogeneity. It has been hypothesized that malignant brain tumor progression and heterogeneity may be governed by selection of specific cancer stem cells (CSCs) over time, due to factors including their genetic, epigenetic and tumor microenvironment (TME) interactions. ^1–4^ The CSCs may leave the tumor bed and acquire additional somatic mutations, ultimately resulting in formation and progression of the tumor in distant areas. As a result of interactions with the TME, some CSCs grow and populate in highly vascular (and therefore, nutrition-sufficient) regions, while others evolve into subpopulations that adapt to more hypoxic (or nutrition-depleted) TME ^5^. Therefore, the co-existence of different subpopulations within the tumor, and their adaptation to the environmental forces, may represent one mechanism for the creation of a robust tumor landscape leading to resistance to standard treatments. In other words, therapies that do not ablate the tumor stem cells will not be effective in eradicating the tumor. Such diversity across the tumor landscape, reflected in radiographic phenotypes of the tumor, allows the tumor cells to adapt to the environmental forces, leading to their clinical behavior and response to therapies.

Glioblastoma (GBM), as defined by the 2021 WHO central nervous system (CNS) tumor classification, is the most common primary malignant brain tumor in adults with a grim prognosis ^6, 7^. GBM tumor cells extensively and diffusely infiltrate throughout the brain parenchyma, rendering it impossible to achieve surgical cure even with maximal safe tumor resection and adjuvant chemoradiotherapy ^8, 9^. In addition, GBM cells form a robust evolutionary ecosystem with multiple genetically-distinct clonal populations, giving rise to intratumor heterogeneity ^10, 11^. It is suggested that standard treatment for GBM (Stupp protocol ^12^), i.e., surgical resection followed by chemoradiotherapy with temozolomide (TMZ), may select resistant subclones from the residual tumor leading to rapid tumor regrowth and treatment failure ^10, 11, 13^.

With recent advances in high throughput sequencing methods, it is now possible to map the genomic alterations and driver mutations in tumors ^14^, which provides opportunities for enrollment in clinical trials or the development of personalized therapies beyond the standard of care for high-grade gliomas ^14^. However, sequencing is generally performed on a small portion of the entire tumor, which, owing to intratumor heterogeneity, is most likely not representative of the whole tumor characteristics ^15^. Multiple mutations can be detected in a single specimen of sequenced tumor. However, it is not straightforward to infer the primary driver genes that lead to the global radiographic phenotype, since tumor cells carrying the mutations are not uniformly distributed across the tumor. Finally, inferences from sequencing depend on the type of the assay, depth of coverage, the comprehensiveness of the libraries, and the implemented analysis techniques, which poses a challenge for implementing uniform pipelines for multi-institutional clinical trials^15^.

Advanced analysis of multiple-sequence MRIs via machine learning methods, often called radiogenomics, has shown great promise for noninvasively probing genomic attributes of the tumor ^16, 17^, while capturing spatial heterogeneity and overcoming sampling bias ^17^. Our present study examines how of the GBM molecular intratumoral heterogeneity impacts imaging features through a comprehensive analysis of its radiogenomics, spatiogenomics (relationship between spatial location patterns of tumor occurrence to mutational composition) and the cross-sectionally estimated molecular evolutionary landscapes. We generate radiogenomic signatures by computational analysis of multiparametric (mpMRI) scans to predict alterations in the driver genes or core signaling pathways based on a targeted sequencing panel. These signatures may provide diverse and complementary information about the underlying properties of the tumor microenvironment (TME), such as neoangiogenesis, cellularity, hypoxia, infiltration. Therefore, they may serve as surrogate *in vivo* biomarkers of the genetic landscape of *de novo* (primary, i.e., Grade 4 tumor at diagnosis) *IDH*-wildtype GBM. We hypothesize that tumors with simpler genomic landscape exhibit more distinctive radiogenomic signatures than heterogenous tumors. We also hypothesize that spatial patterns of tumor locations may be associated with specific molecular signatures or sequence of events, driven partially by characteristics of the cells of origin and partially by the local TME. Finally, we explore the imaging phenotypes that accompany the clonal evolution of tumors.

## 2 Results

### 2.1 Radiogenomic Markers Are More Distinctive in Tumors with Single Gene/Pathway Alterations Compared to Tumors with Multiple Genes/Pathways Altered

Radiogenomic signatures were generated using deep learning (DL) and conventional machine learning (ML) models to predict the presence or absence of alterations, which in this analysis refers to point mutations or insertions/deletions, in the key driver genes and their associated pathways (Supplemental Information, SI1, Table S1). Throughout this section, for brevity, we will refer to a tumor as “mutant”/”wildtype” if it does/does not have alterations in a certain gene or pathway of interest, respectively. For “co-occurring” radiogenomic signatures, the tumors are considered as mutant/wildtype for a gene or pathway if they do/do not have alterations in a gene/pathway of interest, respectively, regardless of the presence or absence of alterations in other genes or pathways.

The cohorts were divided into discovery and independent replication sets to validate the reproducibility of the generated signatures in unseen data. A summary of diagnostic performance of our predictive models in terms of area under the receiver operating characteristic (ROC) curve (AUC) and balanced accuracy, is provided in Supplemental Information, SI4, Table S3. We found high reproducibility of our radiogenomic signatures in independent replication cohorts. DL and support vector machine (SVM) models based only on features computed from conventional MRI scans resulted in comparable distinctive power for most of co-occurring signatures, while the SVM models with features from conventional and advanced MRI achieved better performances. This finding highlights the importance of physiological MRI scans, namely diffusion tensor imaging (DTI) and dynamic susceptibility contrast enhanced (DSC-) perfusion MRI, in characterizing the pathophysiological processes that are derived from molecular alterations in the TME ^18, 19^.

We postulated that radiogenomic signatures become more distinctive in tumors that have mutations in one of the key driver genes/pathways in their mutational landscape, compared to those with mutations in multiple genes/pathways. To test this hypothesis, the distinction of our generated radiogenomic signatures for single-alteration or “exclusive” tumors was investigated. For such radiogenomic signatures, in contrast to the multiple-alterations or “co-occurring” situation, the mutant tumors only had mutations in one gene/pathway of interest, and wildtype tumors had no mutations in any of the genes in the main six pathways (described in the Methods section). The performances of most radiogenomic signatures showed improvement in discrimination of “exclusive” mutant from wildtype tumors (as indicated in Supplemental Information, SI4, Table S3, and Figure S2).

The separability of the signatures for tumors with single alterations, i.e., “exclusive” tumors, is presented in Figure 1(A), as a comparison with the values of the signatures for “co-occurring” tumors with multiple alterations, along with their Cohen’s D effect sizes in Figure 1(B). These results show larger effect sizes for discrimination of tumors with and without mutations for the “exclusive” cases, in contrast with the “co-occurring” tumors for most of the predictive models. This finding suggests that the imaging signatures become clearer and can differentiate the mutant and wildtype tumors better than the tumors with mixed mutation compositions. However, even in the presence of co-occurring mutations, the imaging signatures carry fairly strong distinctive power, as reflected by the effect sizes.

**Figure 1.**
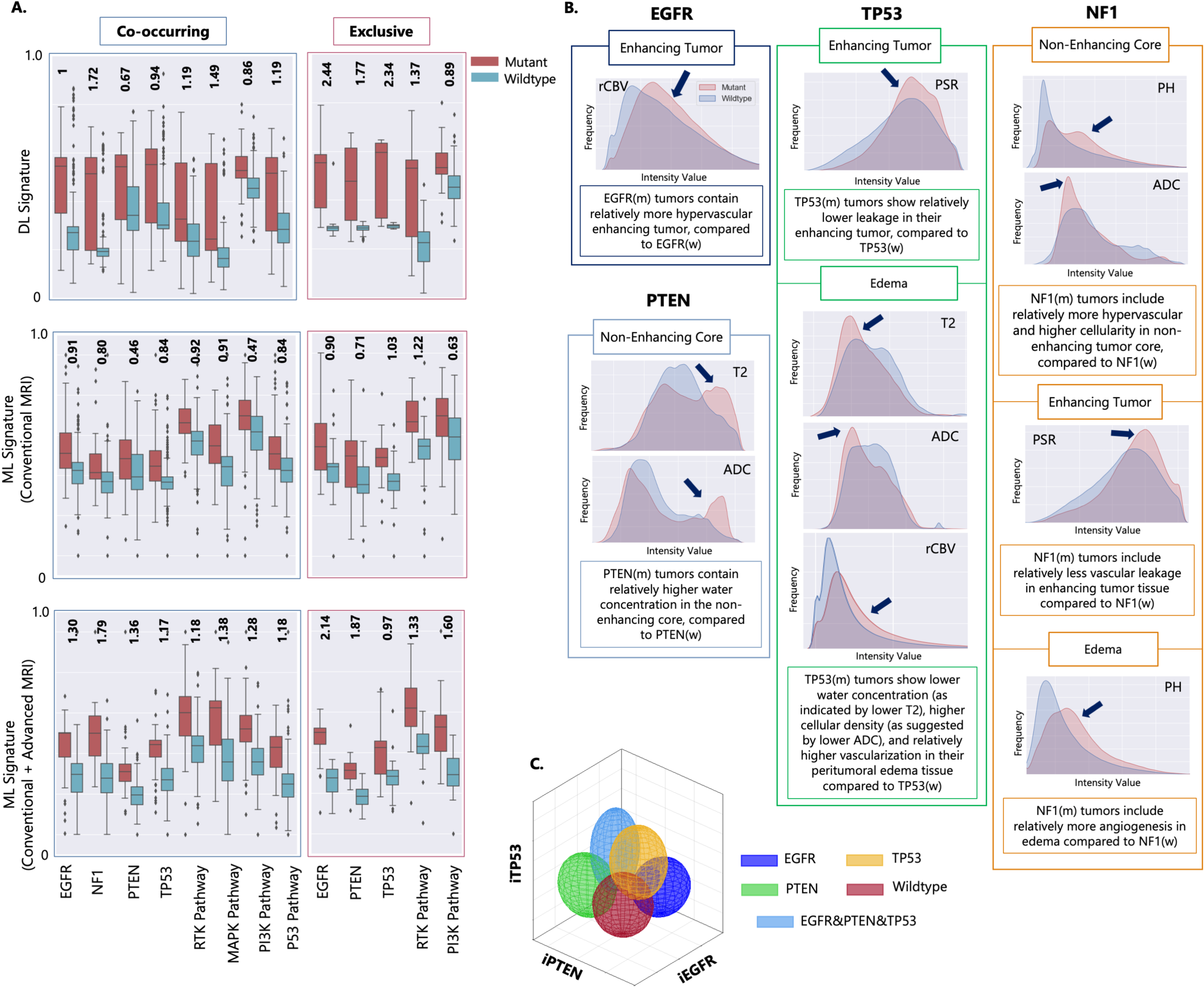
Imaging signatures of IDH-wildtype GBM tumors. (A) Signatures generated using deep learning (DL; top), and conventional machine learning (ML; middle) using only conventional MRI as well as using both conventional and advanced MRI (bottom; details are provided in the Methods section). These signatures in Panel (A) are shown for co-occurring (left) and exclusive (right) mutations and pathways. The numbers indicated at the top of each pair of mutant-wildtype box-whisker plots indicate the effect sizes for the signatures generated by DL, ML (conventional MRI), ML (conventional and advanced MRI) models for co-occurring and mutually exclusive tumors. (B) Representative imaging characteristics in different tumorous subregions for the tumors with mutations in key driver genes in comparison to their wildtype counterparts [rCBV = relative cerebral blood volume; PSR = Percentage Signal Recovery; ADC = Apparent Diffusion Coefficient; PH = Peak Height]. The annotation (m) or (w) after the names of genes, e.g., EGFR(m) or EGFR(w), in the boxes denote mutant or wildtype tumors, respectively. Edema refers to peritumoral edema, which is an infiltrated zone surrounding the tumor comprised of vasogenic edema and infiltrating tumor cells diffused in the adjacent normal brain tissue. (C) A representative example of the degree of expression of imaging signatures, quantitatively measured along respective axes, for mutations in the EGFR, PTEN, and TP53 genes [iEGFR, iPTEN, and iTP53 = imaging signatures of EGFR, PTEN, and TP53 mutations, respectively]. The colors of the ellipsoids represent the molecularly defined mutation status and the shape of the ellipsoids reflects the spread of values of respective imaging signatures. For example, molecularly-defined EGFR mutant tumors (dark blue) tend to express the iEGFR signature, i.e. they have higher values along the iEGFR axis, and so forth.

In the histograms presented in Figure 1(C), the imaging characteristics that are most discriminative between tumors that do and do not have mutations in the various genes are summarized. As the results show, *EGFR* mutant (*EGFR(m)*) tumors exhibit hypervascularity (increased relative cerebral blood volume (rCBV)) in their enhancing tumor region compared to the tumors without *EGFR* changes. Mutations in *TP53* are associated with lower leakage (increased percent signal recovery (PSR)) in the enhancing tumor, decreased water concentration (decreased T2), higher cell density (lower apparent diffusion coefficient (ADC)), and higher neovascularization (elevated rCBV) in their peritumoral edema region. The so-called peritumoral edema manifests as a region of high signal intensity on T2-weighted and FLAIR MRI sequences surrounding the enhancing tumor mass and presents a combination of vasogenic edema and infiltrating glioma cells. Tumors with mutations in *PTEN* show relatively more water concentration (increased T2 signal) and facilitated diffusion (higher ADC) in the non-enhancing tumor region. *NF1*-mutated tumors have relatively more pronounced vascularization (increased peak height (PH)) and reduced diffusion resulting from higher cellularity (lower ADC) in the non-enhancing tumor region. Furthermore, mutations in *NF1* lead to lower vascular leakage (increased PSR) in the enhancing tumor and more angiogenesis (increased PH) in the peritumoral edema.

The scatterplots in Figure 1(D) illustrate the imaging signatures (denoted by *iEGFR, iPTEN,* and *iTP53*) for our ML models created based on conventional and advanced MRI sequences for prediction of mutations in *EGFR, PTEN,* and *TP53* genes. This figure demonstrates the discrimination of tumors for which our sequencing panel identified mutations in a single gene (related to purer molecular landscape) in comparison to tumors with no mutations detected or the tumors with mutations in multiple genes. The values of radiogenomic signatures for the mixed tumors are usually scattered throughout the plot while the values for the tumors with one mutation are more concentrated around a certain region. These findings further confirm that tumors with multiple mutations, or a more heterogeneous molecular landscape, exhibit imaging characteristics which are consistent with the signatures of the co-existing mutations.

### 2.2 Spatiogenomic Landscape of IDH-wildtype GBM: Spatial Distributions Are Associated with Specific Molecular mutations and pathways

To test whether spatial location of the tumor is associated with its mutational profile, statistical atlases capturing the probability that a tumor with certain mutation(s) occurs at a particular brain region (see Methods) were generated. Figure 2 depicts spatial maps of the frequency of occurrence of 358 *IDH*-wildtype GBMs with mutations in each of the key driver genes, i.e., *EGFR*, *PTEN*, *NF1*, and *TP53* (Figure 2(A)), as well as their associated signaling pathways, i.e., *RTK*, *PI3K*, *MAPK*, and *P53* (Figure 2(C)). Furthermore, we generated atlases of spatial patterns for wildtype tumors, i.e., tumors without any mutations in any of the six main pathways, *RTK*, *PI3K*, *MAPK*, *P53*, and *RB1* pathways, and *ChrMod* regulation (including chromatin modifier genes, such as *ATRX*)^20^. Tumors with mutations in certain genes/pathways or their combinations show predilection for specific brain regions, as demonstrated in the atlases on the right side of panels (A) and (C) in Figure 2. We found that *EGFRm* tumors to be more frequent in the left parietal, right temporal, and right occipital lobes, while *PTENm* cases tend to occur more frequently in the left and right temporal, and right parietal lobes (Figure 2A-B). *TP53mt* is more likely to occur in the right temporal and right frontal lobes, and *NF1m* has a predilection for the left frontal lobe (Figure 2A-B). As the results in Figure 2(C) and (D) suggest, subjects with activated *RTK* pathway are more frequent in the right temporal, left parietal, and left frontal lobes, tumors with mutations in the *PI3K* pathway tend to occur in the left and right temporal and left frontal lobes, *P53* activated pathway subjects are more prominent in the right temporal, right frontal lobes, and right basal ganglionic regions, and *MAPK* activated tumors are more likely to grow in the left frontal lobe.

**Figure 2.**
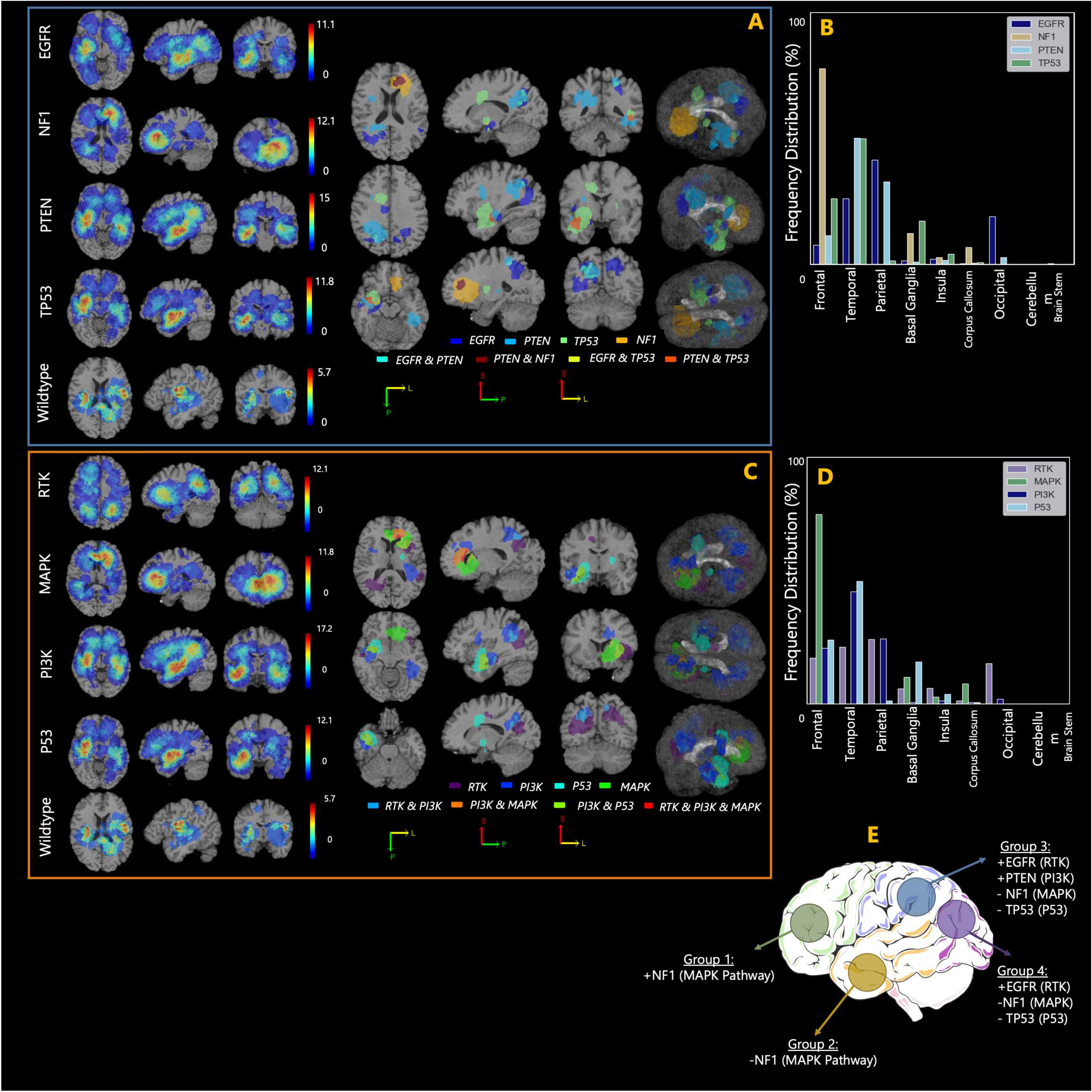
Spatiogenomic landscape of IDH-wildtype GBM. (A) Spatial maps of frequency of occurrence for the tumors with mutations in key driver genes, including EGFR, NF1, PTEN, and TP53, and (C) their associated signaling pathways, i.e., RTK, MAPK, PI3K, and P53. The spatial map of wildtype tumors, i.e., tumors with no mutations in any of the genes involved in the main six signaling pathways (RTK, MAPK, PI3K, P53, RB1, ChrMod) is also shown. In each of subfigures (A) and (C), the maps on the right side indicate the regions with higher likelihood of occurrence of mutations in a gene/pathway or a combination of genes/pathways. The bar plots shown in subfigures (B) and (D) indicate frequency distribution for occurrence of tumors carrying each of the key driver genes/pathways in different brain regions. The basal ganglia label consists of multiple brain regions, i.e., putamen, caudate nucleus, globus pallidus, subthalamic nucleus, nucleus accumbens, internal capsule, and thalamus. All images are displayed in the radiological convention orientation. Subfigure (E) summarizes the findings, illustrating four main groups of regions, towards which the tumors with specific gene mutations show higher predilection.

Based on a population-based approach, four distinct brain regions were derived, representing relatively high prevalence of certain mutations in those regions. A summary of the spatiogenomic markers is provided in Figure 2(E) where four main groups of tumor occurrence propensity are specified. To explore characteristic molecular underpinnings for these groups of tumors we extracted mutational signatures (Figure 3(A)). We used cosine similarity metric (CM) as a measure of closeness among the four mutational profiles (Figure 3(B)). This value ranges from 0 (completely different mutational signatures) to 1 (identical signatures). Mutational signatures were different (CM = 0.35) between Group 1 (frontal lobe) and Group 4 (occipital lobe), which respectively include or lack *NF1* (*MAPK*) mutations. The differences between the spatiogenomic groups were further examined through calculation of mutant-allele tumor heterogeneity (MATH) scores. This score utilizes bulk next-generation sequencing (NGS) data to provide a quantitative measurement of intratumor heterogeneity ^21^. The MATH score is determined by computing the width of the distribution of mutant-allele fractions (MAF) among tumor loci ^21^. In the clonal evolution of the tumor, a locus with earlier mutation has higher MAF and therefore MATH score, while later events result in lower MAF and MATH score ^21^. The Group 4 tumors show significantly higher (p = 0.013) MATH scores compared with the Group 1 tumors.

**Figure 3.**
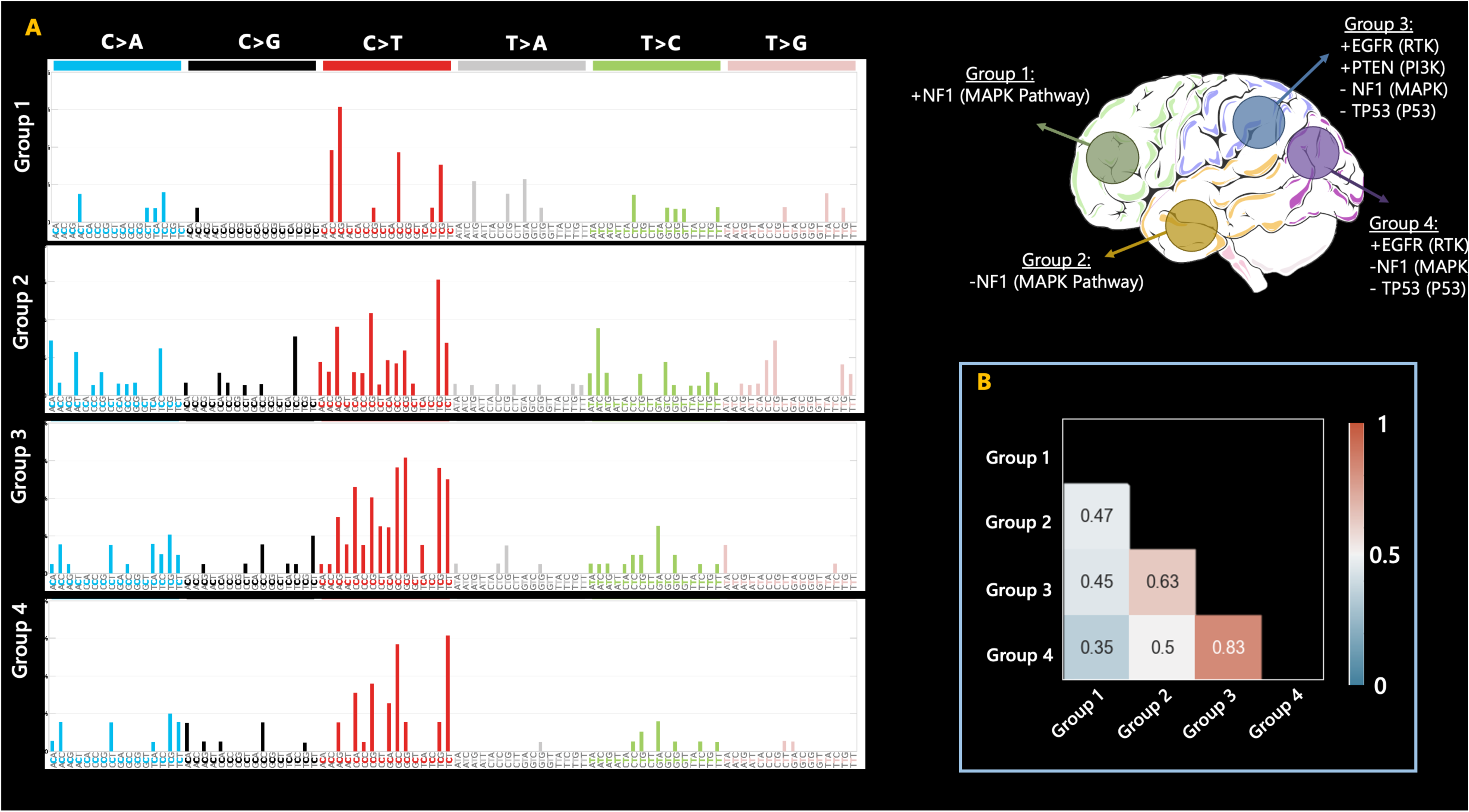
Mutational signatures for the tumors in the four groups of spatial locations. These four groups from Figure 2(E), are illustrated on the top right side of this figure. (A) Mutational signatures; (B) cosine similarity metric (CSM) for the signatures of the four groups, suggestive of least similarity between tumors in groups 1 and 4. MATH scores for the tumors in these four groups showed significant differences in the mean MATH scores between groups 1 and 4 (p = 0.013).

### 2.3 Molecular Underpinnings of Primary IDH-wildtype GBM With Spatial Anatomical Predilection

To elucidate the links between tumor progression or evolution with radiographic appearances, 177 samples with available single nucleotide variants (SNVs) and copy number variation (CN) information were analyzed. Only somatic mutations with MAFs of 5% or greater were included. Figure 3 depicts a summary of common oncogenic mutations, pathway-level alterations, and trajectory evolution model of GBM. The specific details about the statistics of copy number variations in this patient population are provided in Supplemental Information, SI1, Table S2. The high copy number gains and losses were defined as ≥ 5 copies and ≤1 copy, respectively.

The oncoplot (Figure 4(A)) shows that the most prevalent somatic genomic alterations present in these 177 samples were *EGFR* CN gain (54%), *CDKN2A* CN loss (40%) and *PTEN* somatic mutation (39%). These findings are in line with reports that *EGFR* amplification and rearrangement are early events in tumorigenesis ^22^. The pathways with the most frequent genomic alterations were receptor tyrosine kinase (RTK, 74%) pathway, which included *EGFR*, followed by PI3K pathway (54%) which included *PTEN*, followed by RB1 pathway (48%) including *CDKN2A* (Fig 4 (B)). Overall, more than 90% of the tumors have RTK and/or PI3K alterations (Fig 4 (B)). The circos plot (Figure 4(D)) shows that the most co-mutated alterations were *EGFR* snv – *EGFR* amplification, *EGFR* amplification – *CDKN2A* deletion, and *EGFR* amplification – *PTEN* snv.

**Figure 4.**
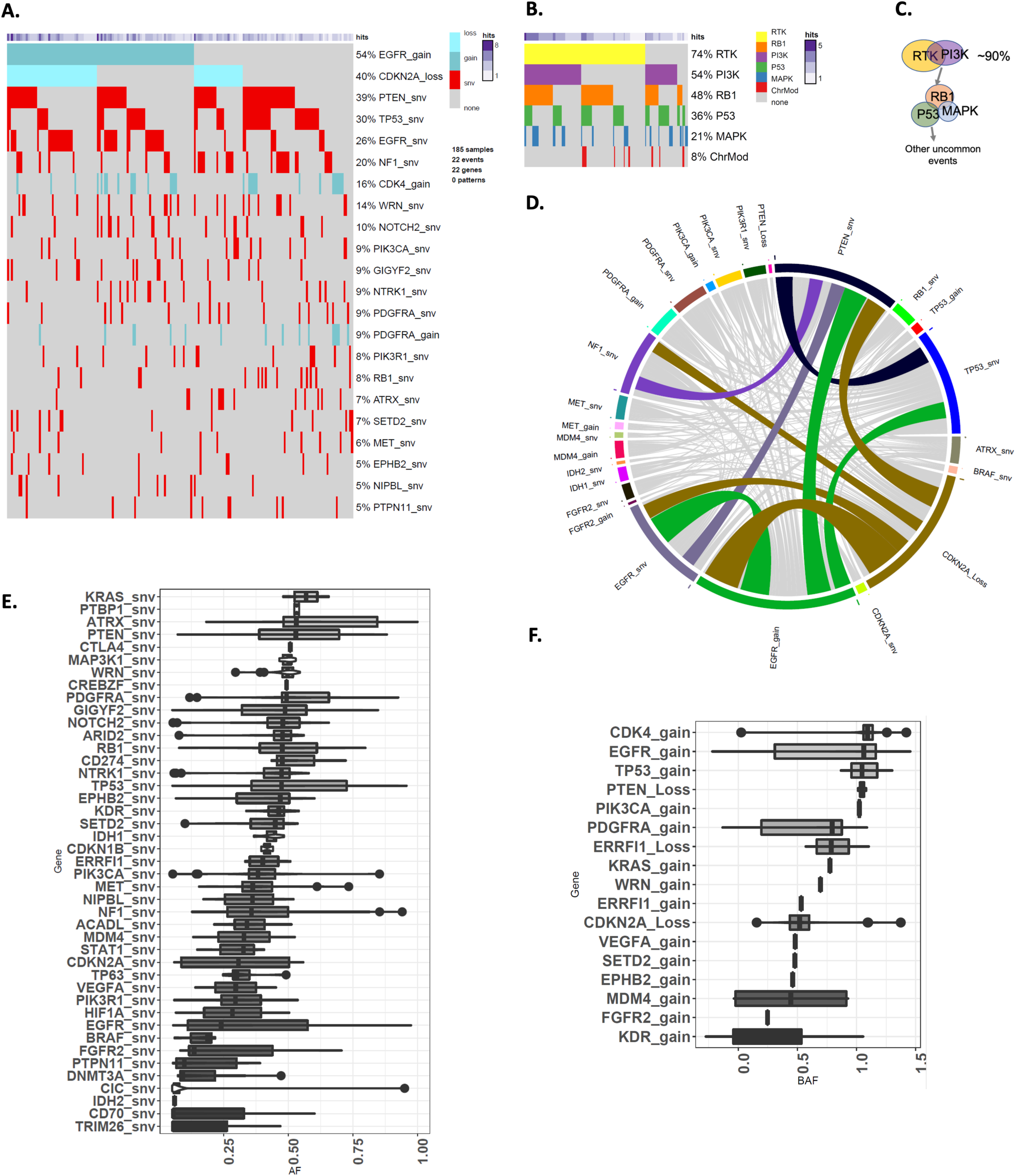
A summary of common oncogenic mutations, pathway-level alterations, and trajectory evolution model of GBM. (A) Heatmap showing the somatic mutations including single nucleotide variant (snv) and copy number changes identified from target exon sequencing of 177 tumors. Snv, CN deletion and amplification are separately denoted by red, blue and dark blue color. (B) Heatmap showing the six mutation pathways including 35 mutated genes in 177 tumor samples. The bar graph on the top depicts the number of tumors. (C) Schematic representation of evolutionary trajectory model of GBM progression that shows the estimated order of events at the pathway level. (D) Circos plot showing the co-mutated genes shared by tumors; colored connections present co- mutated genes shared by more than 20% of tumors and the grey color connections denote co-mutated genes shared by less than 20% of samples. The width of the connection represents the sample size. (E) The allele frequency for all sequenced genes, (F) B allele frequency for genes with CN alterations (gains and losses).

Next, we examined whether the tumors differ in their order of events of somatic genomic alterations during GBM evolution using a Bayesian inference approach implemented in TRONCO^23^. In brief, this approach combines techniques for sample stratification, driver selection, and identification of fitness-equivalent exclusive alterations to predict likely temporal order of mutational events during the course of disease progression based on Suppes’s probabilistic causation. The trajectory of GBM progression was inferred based on mutated genes included in the six signaling pathways that were present in more than 10% of all samples. The model captured the early drivers including *EGFR* amplification, and somatic mutations in *NF1*, *PTEN* and *TP53* (Supplemental Information, SI5, Figure S3). *EGFR* amplification was accompanied by elevated allele frequency and mutations in *PTEN* and *TP53* also showed higher allele frequencies (Figure 4 (E-F)). *EGFR* point mutations were detectable at a moderately low allele frequency (Figure 4 (E)).

Next, we postulated that the differences in oncogenic alterations and evolutionary trajectory are manifested in tumor phenotypes, ultimately reflected in radiogenomic features and therefore investigated the evolution patterns for the tumors originating in the distinctive brain regions of Groups1-4 of Figure 2.

Figure 5 provides the evolutionary trajectories for tumors in the four brain regions shown in the middle (regions of relatively distinct molecular profiles). As suggested by the results, *NF1* alterations (100%) were identified as the most frequent and potentially early event in Group 1. In contrast, *PTEN* mutations (56%), *EGFR* amplification (56%), and *EGFR* mutations (44%) were the most common events in Group 2, followed by *TP53* (38%) and *NOTCH2* (31%) mutations. *PTEN* mutation and *EGFR* amplification were the likely early and driver mutations in Group 2 tumors, but it was not possible to ascertain the preferential precedence between those two with statistical significance. In Group 3, *PTEN* mutations (72%), *EGFR* amplification (61%), and *EGFR* mutations (33%) were the most frequent events, and the evolutionary analysis identified *PTEN* mutation and *EGFR* amplification as the likely early driver events. In addition, *TP53* and *CDKN2A* mutations were also captured as additional driver mutations. The most frequently occurring alterations in Group 4 were *EGFR* mutations (86%), *EGFR* amplification (71%) and *GIGY2* mutations (43%). The evolutionary trajectories also indicated the possibility of *EGFR* and *PTEN* mutations as early events in the progression of tumor growth, but sample sizes were small and inferences did not reach significance level (*p*>0.05). The derived evolutionary trajectories for all the four groups are provided in more detail in (Supplemental Information, SI5, Figures S4-S7).

**Figure 5.**
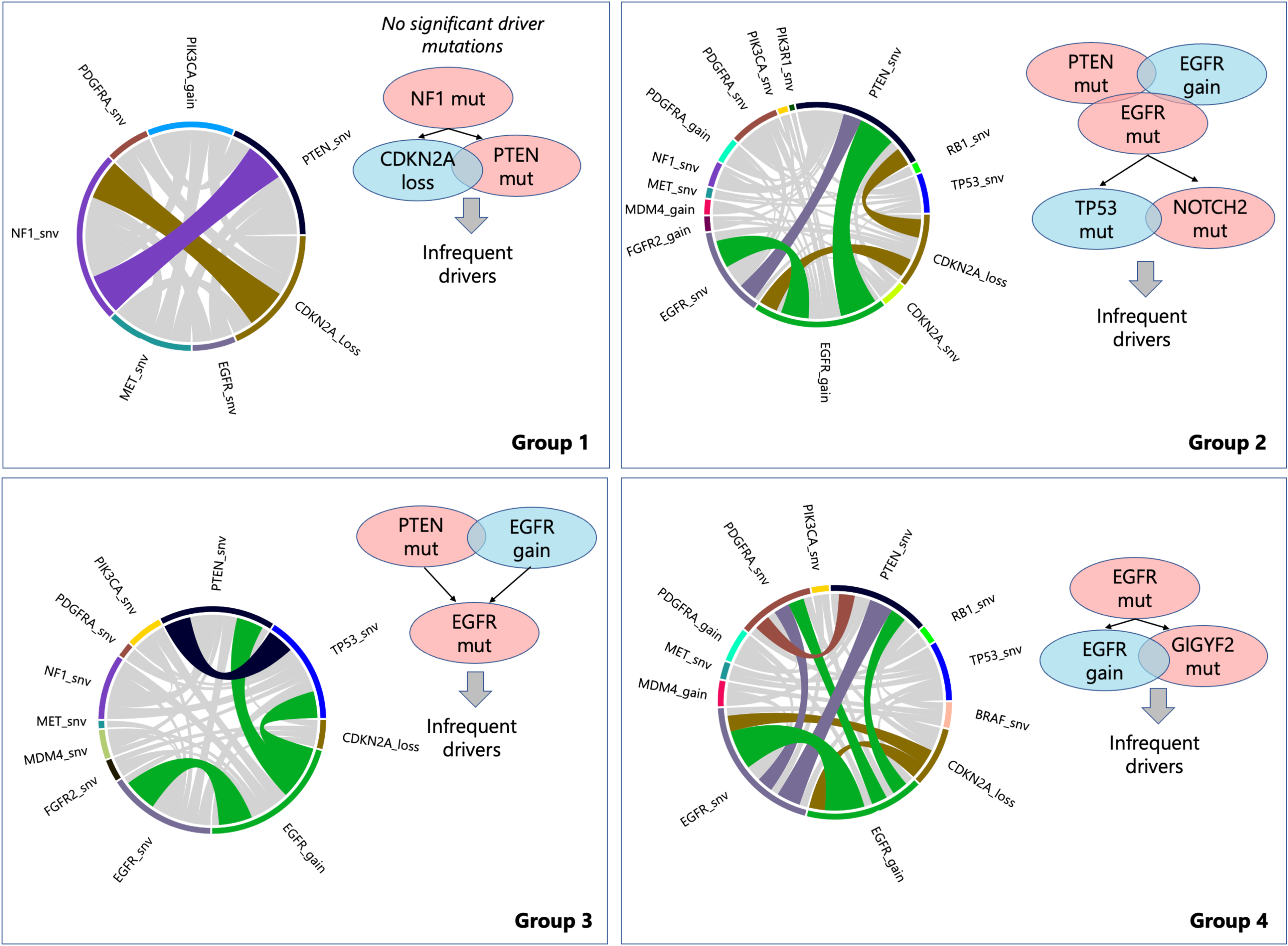
Trajectory evolution model for the four found brain region. Schematic representation showing a summary of the order of events in GBM progression. *NF1* alterations were identified as the most frequent and potentially early event in Group 1. In Group 2, *PTEN* mutations, *EGFR* amplification, and *EGFR* mutations were the most common events, followed by *TP53* and *NOTCH2* mutations. In Group 3, *PTEN* mutations, *EGFR* amplification, and *EGFR* mutations were the most frequent events. In Group 4, *EGFR* mutations, *EGFR* amplification and *GIGY2* mutations were most frequent alterations.

## 3 Discussion

Cancer arises as a result of genetic and epigenetic alterations in primary founding clones^24^. However, by the time the tumor is detected, the tumor population size has significantly expanded with presence of subclones representing intratumor heterogeneity ^24^. Malignant cells usually carry somatic mutations that were present in their founder clone along with additional mutations they acquired during tumor progression ^24^. Like other human cancers, formation and progression of GBM is accompanied with activation of oncogenes, inactivation of tumor suppressor genes, deregulation of DNA repair genes, genomic copy number changes, and vast epigenetic alterations ^25^. In GBM, such alterations can occur in different subpopulations that cohabit within the same tumor, giving rise to intratumor heterogeneity, thereby, triggering appearance of different imaging characteristics. In our study, we provided insights into the inter-connections between the neuroimaging patterns or radiophenotypes and the underlying molecular underpinnings of *IDH*-wildtype GBMs, thereby elucidating different mechanisms by which various mutations influence the phenotype of the tumor and its surrounding infiltrated brain tissue. We attempted to bridge the gap between the information from genomic and imaging scales by mapping the molecular landscape of *IDH*-wildtype GBMs using spatial pattern analysis (spatiogenomic) and multivariate quantification of clinically obtained mpMRI scans.

### 3.1 Imaging – Genomics Landscape of GBM

Our findings provide evidence for several associations between *in vivo* imaging characteristics in GBM and underlying genetics: (1) discovery of non-invasive and highly distinctive radiogenomic signatures of gene or pathway alterations, (2) the impact of molecular heterogeneity on distinctiveness of the derived imaging signatures, (3) associations between spatial location of tumors and their molecular underpinnings, as suggested by cross-sectionally estimated evolutionary trajectories. Previous studies on GBM radiogenomics ^18, 19, 26–29^ were limited to exploring correlations of specific gene mutations with imaging phenotypes or generating predictive radiomic models for single targets without assessment of other simultaneously co-occurring mutations. Multiple studies have attempted to use mpMRI of the whole tumor and tumor subregions, including enhancing tumor, nonenhancing tumor core, and peritumoral edema, to differentiate radiogenomic signatures of individual genes ^18, 19, 26–29^. However, an integrative and extensive study of several genes and pathways has been missing from former studies. Herein, we found that a comprehensive radiogenomics panel can be derived using AI methods, thereby providing noninvasive assessment of the degree of phenotypic expression of alterations in the driver genes/pathways on an individual patient basis. Furthermore, this is the first study of its kind to fully explore the molecular landscape of GBM and its alterations that are manifested in the imaging features, thereby offering mechanistic insights into how various mutations shape the phenotype of the tumor and its surrounding infiltrated brain tissue.

#### 3.1.1 Molecular Heterogeneity

Our results demonstrated that molecular heterogeneity of tumors can be captured by their radio-phenotypes. Identifying the genomic variations in the tumor (or tumor genotype) helps gain better understanding of the tumor ecosystem. However, tumor evolution is largely driven by the interactions of genetically defined phenotypic traits with the environmental selection forces. Therefore, comprehensive characterization of the tumor, including its molecular, spatial and temporal heterogeneity, can be better assessed with the availability of *in vivo* phenotypic biomarkers extracted from radiographic images ^30^. We elucidated the association between phenotypic variations that result from genotypic alterations by providing evidence that radiogenomic signatures become more distinctive in tumors that are molecularly less heterogeneous, compared with tumors with higher intratumor molecular heterogeneity. This *in vivo* estimation of molecular heterogeneity and the degree of phenotypic expression of different genetic alterations in a tumor can inform patient management and clinical trial stratification, especially in molecularly driven therapies.

#### 3.1.2 Imaging Characteristics offer Mechanistic Insights

Our results uncovered many imaging characteristics specific to tumors bearing mutations in driver genes and pathways, which offer mechanistic insights into how mutations shape the phenotype of the tumor and its surrounding infiltrated brain region, including neo- vascularization, cell density, and peri-tumoral infiltration and migration. Tumors with mutations in *EGFR* were found to be associated with features indicative of higher angiogenesis in the active (enhancing) tumor region, as compared to the tumors without *EGFR* mutations. This finding is in agreement with the involvement of amplified and/or overexpressed growth factors, or oncogenes, including *PDGF*, *EGFR*, *MET*, and *FGFR*, in overgrowth of high-grade gliomas ^31^. The *EGFR* gene, as one of the most frequently altered genes in the *RTK* pathway, encodes the EGFR protein, which is a transmembrane tyrosine kinase receptor, and is critical in regulating cell division and death. Ligand-receptor binding transmits signals that result in activation of a series of intracellular signaling cascades affecting apoptosis, angiogenesis, and invasion ^31^. Alterations in *EGFR* can lead to constitutive activation of the signaling pathway ^31^, and the resulting phenotype is detectable by our imaging analysis.

*PTEN*, in the *PI3K*-AKT-mTOR signaling pathway, is a tumor suppressor gene that regulates cell division to prevent uncontrolled or rapid proliferation, inhibits transmission of growth factor signals through the *PI3K/AKT* signaling pathway, triggers the apoptosis process, and is involved in cell migration, angiogenesis, and adhesion of cells to the surrounding tissue ^32^. *PTEN* loss or downregulation is an early phenomenon and among the most frequent mutations in GBM ^17^. Our findings on the association of *PTEN* mutations with relatively less dense tissue in the non-enhancing core (including necrotic areas), as suggested by increased T2 signal and ADC values, aligns with the reported role of elevated water content in the necrotic core in facilitating invasive tumor growth and its penetration in the surrounding tissue ^34^ and involvement of *PTEN* mutations in necrogenesis and tumor invasiveness ^35^ . Increased water density can reflect lower tumor density, potentially due to pre-necrotic processes.

*TP53* maps to chromosome *17p* and encodes the p53 protein, which has an integral role in cell cycle arrest, response to DNA damage, and apoptosis; therefore, its inactivation leads to tumor progression through genomic instability ^37^. In our study, mutations in the *p53* pathway are accompanied by elevated cellular density (lower ADC), decreased water concentration (lower T2), and relatively increased perfusion (higher rCBV) in peritumoral edema, suggestive of diffuse infiltration. Given that the inactivation of genes in p53 pathway regulate the cell cycle by governing the G1-to-S phase transition ^36^, alterations in this cell cycle regulator expose the tumors to abundant cell division via mitogenic signaling effectors, including *PI3K* and *MAPK* ^36^, with one potential downstream effect being expansion of the tumor by infiltration further into the brain, as suggested by imaging *NF1* is a tumor suppressor gene that encodes neurofibromin protein, a negative regulator of cell growth through suppression of RAS activation ^38^. Therefore, losses in neurofibromin functionality, as a result of inactivating mutations in *NF1*, will prolong activation of the RAS/MAPK signaling pathway, cause a loss in growth control and increase cellular proliferation ^39^. In addition, reduced expression of neurofibromin confers resistance to therapeutic drugs, a fact that highlights the importance of detecting alterations in this pathway clinically ^39^. *NF1* alterations and RAS/MAPK pathway activation have been reported to prevail in a majority of mesenchymal subtype GBMs, which have the lowest survival rates among GBM molecular subtypes ^20, 40^. Such tumors exhibit a high degree of macrophage/microglia infiltration, express high levels of angiogenic markers and demonstrate extensive necrosis ^41, 42^. These biological characteristics of *NF1* mutations supports our findings of relatively greater vascularization and increased cellular density within the non-enhancing tumor core (that also contains pre-necrotic regions) and higher angiogenesis in peritumoral edema in tumors with presence of mutations in *NF1*.

Radiogenomic signatures in the tumors with mutations in genes/pathways of interest generally showed higher variances compared to their wildtype counterparts. As our multivariate analysis approach focused on capturing heterogeneity within each tumor subregion, quantified by radiomic features, this finding is indicative of higher heterogeneity within the mutated tumors compared to the wildtypes ^19^. This highlights the role of the proposed radiogenomic signatures in complementing the findings obtained via molecular analysis by overcoming the limitations posed by sampling bias in such a heterogeneous tumor ^19^.

As presented in our study, in addition to imaging features of the central core area that are indicative of tumor genotype, imaging features derived from peritumoral edema contributed to distinction of the tumors with or without mutations in various genes (e.g., mutations in *TP53* and *NF1* genes that accompanied alterations in water content, infiltration, and neo-angiogenesis). This tumor subregion is formed as a biological response to diffuse infiltration of glioma cells into normal tissue surrounding the tumor, which causes the release of angiogenic and vascular permeability factors ^27^. As resection of peritumoral edema, the propagating front of the tumor, is not usually a part of standard of care treatment for GBMs, future tumor recurrence becomes inevitable. Better understanding of radiogenomic characteristics of peritumoral edema can potentially aid in exploiting alternative local therapies such as extended resection, intensified radiation, focused ultrasound, intra-operative viral injections, or convection enhanced intratumoral delivery of therapeutic agents.

### 3.2 Spatial Pattern – Genomics Landscape

Our study demonstrated a spatial predilection of tumors with mutations in key driver genes, consistent with findings of related lower-scale studies ^43, 44^. Critically, our findings elucidate the differences in molecular underpinnings of tumors occurring in different brain regions. The spatial distribution of the tumors in our study may indicate the contribution of region-specific precursor cells. We found clusters of tumors in the perisylvian regions, as well as in subcortical white matter with extension to superficial grey matter, which aligns with the previous findings about presence of neural CSCs in regions of the brain, such as the dentate gyrus ^44^. GBMs likely arise from neural progenitor cells or stem cells. Astrocyte-like neural CSCs in the subventricular zone with low-level somatic mutations are the likely origin of *IDH*-wildtype GBM ^45^. We show that molecular heterogeneity, to the extent that can be measured by our targeted sequencing panel and MATH, and the similarity of the mutational signature for the group of tumors located in the frontal lobe are statistically lower (p<0.05) than heterogeneity and signatures of the tumors clustered in the occipital lobe, and lower than the clusters of tumors in the temporal and parietal lobes (although not found statistically significant, p>0.05). This distinction in the molecular composition of the tumors in the discovered tumor region groups was also observed by analyzing the evolutionary trajectories, which were estimated via statistical modeling of the pre-surgical genomic data. As the findings suggest, the tumors in the cluster concentrated around the frontal lobe may undertake a different progression path compared to tumors that occur in other regions, especially those arising in proximity to the occipital lobe.

It is proposed in the literature that tumor progression and heterogeneity are governed by selection of specific CSCs over time, due to factors including their genetic, epigenetic and TME interactions ^1–4^. The progenitor CSCs may migrate from their origin and acquire additional somatic mutations, ultimately resulting in formation and progression of the tumor in distant brain parenchyma. Interactions with the TME can give rise to growth of some CSCs in highly vascular regions, or adaptation of some subpopulations to more hypoxic TME ^5^. Numerous studies have reported on the role of the TME in forming the tumor phenotype and giving rise to its spatial and temporal heterogeneity ^46^. The distinction in spatiogenomic signatures of tumors located in different brain regions, as supported by their modeled temporal evolution path, further highlights the contribution of TME in building the underlying molecular landscape of GBM.

### 3.3 Limitations

Our study has limitations. While the sample size for this study was larger than several other radiogenomic studies of GBM, it would benefit from a larger cohort, in which additional pathway level alterations can be detected and be correlated to the radiogenomic and spatiogenomic characteristics. Limited sampling of biospecimen for genomic profiling and targeted gene panel sequencing may not capture certain classes of genomic alterations (e.g. fusions) and underappreciate intratumor genetic heterogeneity in driver mutations. Formalin fixation of the tumor samples that is carried out for diagnostic tests in molecular pathology can cause DNA damage, such as fragmentation and non-reproducible sequencing artifacts ^47^. In our analysis of mutational signatures, disproportionate levels of C>T changes that appeared in all our four signatures may have been a result of these artifacts. However, as we used cosine similarity metric to determine the distinct patterns of mutational signatures, we do not believe that these artifacts that were similar among all four signatures influenced the derived conclusions.

### 3.4 Conclusion

In conclusion, we have elucidated the links between regional variations expressed as radiophenotypes and the molecular landscape of *IDH*-wildtype GBM. Prompted by the growing advances in computational analytics in the radiology – neuro-oncology domain and evidence for the potential of imaging to reveal molecular characteristics of the tumors, we comprehensively analyzed the radiogenomic, spatiogenomic, and evolutionary landscapes of GBMs to provide better understanding of the links between radio- phenotypes and the underlying molecular heterogeneity of GBMs. We found distinct radiogenomic signatures of mutations in several genes and signaling pathways that became more pronounced when the tumors were less molecularly heterogeneous. Importanty, these signatures offered mechanistic insights into how various mutations shape the phenotype of the tumor and its surrounding brain tissue via changes in cell density and neo-vascularization. We further discovered spatiogenomic signatures from tumors that carry certain mutations, suggesting the association of molecular composition with the brain regions they arise from. We showed that these spatial patterns may emerge as a result of differences in TMEs in different brain regions that give rise to distinct spatial and temporal heterogeneity and define molecular composition of the tumors. The findings of this study can assist in noninvasive identification of patients who may benefit the most from molecular targeted therapies, as well as allow for longitudinal monitoring of mutational changes during treatment.

## 4 Methods

### 4.1 Study Cohort

This HIPAA-compliant study was approved by the IRB. All patients provided their informed consent at the time of imaging. A retrospective cohort of n = 585 patients with GBM tumors who had undergone surgery at the Hospital of the University of Pennsylvania (HUP) between 2006 to 2019 was reviewed. The patients who had undergone maximal safe or partial resection, were histopathologically confirmed with *de novo* glioblastoma tumors, received radiotherapy, and concomitant and adjuvant chemotherapy with temozolomide, and at the very least had four conventional MRI scans, including pre- and post-gadolinium T1-weighted (T1, and T1-Gd), T2-weighted (T2), and T2 fluid attenuation inversion recovery (T2-FLAIR), acquired pre-operatively on a 3T MRI scanner (Siemens, Tim Trio, Erlangen, Germany) for a first-occurrence brain tumor were included. A cohort of n = 374 patients who met the inclusion criteria was collected.

### 4.2 Next Generation Sequencing (NGS)

#### 4.2.1 NGS Panel 1

Sequencing was performed for n = 183 subjects using an in-house research NGS panel. A custom AmpliSeq sequencing panel was designed to target the coding sequence of 45 genes and 4 additional mutation loci of interest using the Ion AmpliSeq Designer portal (ThermoFisher, https://ampliseq.com/). The design consists of 2130 amplicons covering 206 kbp, with amplicon size range optimized for FFPE DNA (125bp - 175bp). Barcoded libraries were prepared from nanogram quantities of FFPE gDNA using the Ion AmpliSeq Library Kit 2.0 (ThermoFisher cat. no. 4480442) as described by the manufacturer. Libraries were quantified using the Ion Library TaqMan Quantitation Kit (cat. no. 4468802) eliminating the need for amplification of the final library. Library concentrations were normalized and pooled in equimolar ratios for multiplex sequencing on an Ion Torrent S5 sequencing system using the Ion 540 Kit-OT2 and Ion 540 Chip (cat. nos. A27753 and A27766).

Signal processing and base calling, followed by alignment of sequence reads to the hg19 reference genome, were performed on instrument using Torrent Suite Software version 5.8. Library performance was assessed using the Torrent Suite coverage analysis plugin. Aligned reads in BAM format were uploaded to the cloud-based Ion Reporter Software (ThermoFisher, https://ionreporter.thermofisher.com/) for variant calling using the version 5.10 default single sample somatic variant calling parameters. Ion Reporter analyses were downloaded to facilitate further processing of the resulting VCF files.

Variant calls were normalized, multiallelic sites split, and NOCALL or reference calls removed using BCFtools. Normalized variants were annotated with Annovar then filtered on variant quality metrics and annotations, again with BCFtools. Variants with variant quality score < 30, read depth < 100, allele frequency < 0.05, number of alternate reads < 5, strand bias > 0.7, as well indels called in a homopolymer greater than 8bp in length, were removed. Also removed were any variants identified as likely artifacts by (1) low population frequency (gnomAD exome or genome frequency < 1%) AND (2) presence in more than 2 of 10 HapMap samples obtained from the Coriell Institute and sequenced by the same means. The 10 control samples include the following DNA sample obtained from the NIGMS Human Genetic Cell Repository at the Coriell Institute for Medical Research: NA12342. Annotation filters retained only exonic and splicing variants with gnomAD exome or genome population frequency less than 1%.

Target genes in this sequencing panel included TP53, PTEN, ATRX, EGFR, VEGF, PDGFRA, PIK3CA, PIK3R1, NF1, PDL1, CTLA4, HIF1A, MDM4, RB1, STAT1, CD70, CIC, FUBP1, CDK4, ACADL, TRIM26, SMAD1, ARID2, CDKN1B, CREBZF, DNMT3A, EPHB2, ERRFI1, FGFR2, GIGYF2, KDR, KRAS, MAP3K1, MET, NIPBL, NOTCH2, NRAS, NTRK1, PTBP1, PTPN11, SETD2, SMARCB1, TP63, WRN. The hotspots included IDH1 p.R132, IDH2 p.R172, BRAF p.V600, H3F3A p.K28, TERT (chr5:1295151-1295315). H3F3A and TERT were excluded, the former was not effectively covered due to presence of pseudogene that was ignored by the variant caller, and the latter had poor amplicon performance.

#### 4.2.2 NGS Panel 2

Genetic data was available for n = 191 patients through a clinical sequencing panel of 153 actionable and prognostic genes for solid tumors, implemented and validated at our institution for clinical assessment of the resected tumors. The panel has a full coverage of all included genes with its Agilent Haloplex design. Full description of this panel and the in-house data processing bioinformatics pipeline can be found in ^48^.

#### 4.2.3 Final Mutational Data

A total of 27 genes in both panels were included for n = 374 subjects and used for our radiogenomic analysis: *ARID2*, *ATRX*, *BRAF*, *CDKN2A*, *CIC*, *DNMT3A*, *EGFR*, *FGFR2*, *FUBP1*, *IDH1*, *IDH2*, *KDR*, *KRAS*, *MDM4*, *MET*, *NF1*, *NOTCH2*, *NTRK1*, *PDGFRA*, *PIK3CA*, *PIK3R1*, *PTEN*, *PTPN11*, *RB1*, *SETD2*, *SMARCB1*, *TP53*. We excluded patients with mutations in *IDH1* or *IDH2* (n = 16). Finally, a cohort of n = 358 patients with *IDH*-wildtype GBM (a summary of demographics, clinical, and genomic characteristics can be found in Supplemental Information, SI1, **Table S1**) was considered for building the population atlases of spatial locations and generating radiogenomics signatures based on conventional MRI (T1, T1-Gd, T2, and FLAIR) scans, of which a subset of n = 228 patients had advanced imaging scans (DTI and DSC-MRI) along with conventional MRI.

We considered the GBM pathways introduced in the seminal paper by Brennan *et al* ^20^, including RTK pathway (with *EGFR*, *PDGFRA*, *MET*, *FGFR2* genes), PI3K pathway (including *PIK3CA*, *PIK3R1*, *PTEN* genes), MAPK pathway (*NF1*, *BRAF*), P53 pathway (*TP53*, *MDM4*), RB1 pathway (*CDKN2A*, *RB1*), and ChrMod pathway (*ATRX*).

### 4.3 Mutational Signature Analysis

We extracted mutational signatures to explore characteristic molecular underpinnings for the groups of patients with specific spatial predisposition. The analysis was performed in accordance with the method proposed by Alexandrov *et al* ^49^. *De novo* global-signature discovery was performed using sigProfilerExtractor ^49^ with 100 iterations to extract each signature and 96 trinucleotide sequence contexts were included. Furthermore, cosine similarity metric was calculated between the obtained signatures for each group of tumors to quantify and validate different mutational profiles between the groups.

### 4.4 MATH Score

We imported the clinical data and NGS sequencing results for 183 GBMs. MATH score for each tumor was calculated from the median absolute deviation (MAD) and the median of its mutant-allele fractions at tumor-specific mutated loci according to the following formula ^21^:

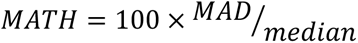

### 4.5 Tumor Evolutionary Trajectories

#### 4.5.1 Analysis of Copy Number Variation

CNVs were called from the Ion Torrent sequence read alignments (BAM files) using CNVkit (version 0.9.9) ^50^. An FFPE CNV baseline was bootstrapped using ten HapMap control samples obtained from Coriell and FFPE tissue derived DNA from 14 lung metastases to the brain, all sequenced with the AmpliSeq panel. The HapMap samples were screened for the absence of CNVs in panel genes in DGV, but an FFPE baseline is desirable to more accurately represent the variability in sequencing coverage expected from the cases. An initial baseline was generated using the HapMap samples, against which the lung metastases were analyzed. The lung metastases were then filtered based on CNV calls to retain 6 samples for which all calls were either single copy gain or loss, and no two samples contained a call in the same gene. These six samples were used to generate an FFPE CNV baseline for analysis of the AmpliSeq cohort.

The case and control sample BAM files were first processed using CNVkit’s ’batch’ command in ’targeted amplicon sequencing’ mode with the ’--drop-low-coverage’ flag, using the UCSC hg19 refFlat.txt file for gene annotation. CNVkit splits the targeted regions into bins and calculates coverage tables for each sample. A pooled copy number reference is generated from the control samples’ coverage tables then used to correct case sample coverage for regional biases and calculate normalized copy number ratios. CNVkit’s ’segment’ command uses the ’circular binary’ method to generate copy number segments from bins. Because the 45 panel genes are sparsely distributed across 18 chromosomes, this approach resulted in segments combining genes separated by large genomic distances. To avoid this behavior bins were instead segmented per gene using the ’genemetrics’ command. Segments consisting of less than 5 bins were dropped, and a copy number ratio threshold of 0 was used to prevent copy number calls at this stage. Segments were then annotated with additional summary statistics using the ’segmetrics’ command to support downstream QC and filtering.

Using the ’call’ command, copy number ratios were adjusted for tumor cellularity and converted to integer copy number values using the ’threshold’ method. Copy number ratio thresholds of -1 and 0.58 were selected to correspond to strict copy loss and gain respectively ^51, 52^. Germline variant allele frequencies were input during the ’call’ step resulting in output annotated with the average b-allele frequencies of heterozygous SNVs in each segment. Finally, CNVs were filtered using a custom script to remove calls for which the log2 ratio 95% confidence interval contains 0, the balanced case.

The default for CNVkit is to calculate log2 ratios and copy number with respect to a diploid chrX and haploid chrY, infer sample gender based on chrX and chrY coverage, and double chrX coverage for male samples. Because this panel targets only one gene on chrX and none on chrY, gender inference was not always accurate. Additionally, the ’genemetrics’ and ’call’ commands are not typically used in conjunction, and both attempt to determine gender, which can result in unexpected behavior. To address this all samples were treated as female at the ’genemetrics’ step while gender was explicitly asserted in the ’call’ command.

#### 4.5.2 Prediction of Evolutionary Trajectories

The oncoprint function implemented in maftools package ^53^ was used for visualization of somatic mutations and pathway level alterations in the GBM samples. We performed the evolutionary analysis using the CAPRI algorithm implemented in TRONCO ^23^ with its default parameters. Both Akaike and Bayesian information criteria were used for regularization to prevent overfitting. The evolutionary trajectories were drawn using sem- supervised approach to improve clarity and interpretability. We used chordDiagram function implemented in circlize package ^54^ to visualize the co-mutated genes in GBM tumor samples. ggplot2 package ^55^ was used to visualize the allele frequency for all mutate genes. All analysis was conducted in R 3.6.0 (2019-04-26).

### 4.6 Radiogenomic Signatures

#### 4.6.1 Image Pre-Processing

Pre-processing of multi-parametric MRI (mpMRI) scans was performed using the CaPTk software (https://www.med.upenn.edu/cbica/captk/) with the following steps: (1) conversion of MRI volumes from DICOM to Neuroimaging Informatics Technology Initiative (NIfTI) format; (2) re-orientation to a reference coordinate system (here, left-posterior-superior (LPS)); (3) co-registration and resampling to an isotropic resolution of 1mm^3^ based on a common anatomical SRI24 atlas ^56^, performed using the “Greedy” software ^57^; (4) skull-stripping of the MRI volumes using the DeepMedic software ^58^, followed by manual revisions where necessary.

Automatic segmentation of tumorous subregions, i.e., enhancing tumor (ET), nonenhancing tumor core (NC), and peritumoral edema (ED), was applied using DeepMedic followed by manual revisions where necessary. An additional segmentation label, namely the whole tumor (WT=ET+NC+ED), was produced by taking the union of all three subregions. Finally, the image intensities were rescaled in the intensity range of 0-255 after removing outliers.

For radiogenomics signatures, in addition to the structural images described above, quantification of DTI scans was performed to generate maps of axial diffusivity (DTI-AD), trace (DTI-TR), radial diffusivity (DTI-RD), and fractional anisotropy (DTI-FA), and DSC-MRI perfusion scans to create maps of peak height (DSC-PH), percentage of signal recovery (DSC-PSR), and an automatically extracted proxy to relative cerebral blood volume (DSC-rCBV). For DSC-MRI, principal component analysis (PCA) was performed to reduce the dimensionality of perfusion time series to 5 components that represent the temporal dynamics of tissue perfusion. More details about quantification of perfusion PCA can be found in ^59^. As a result, voxel-wise parametric maps of five PCAs (DSC-PC1, DSC- PC2, DSC-PC3, DSC-PC4, and DSC-PC5) were generated.

#### 4.6.1 Atlases of Spatial Location

We applied the Glioma Image Segmentation and Registration (GLISTR) software ^60^ to generate population atlases of spatial distributions. GLISTR simultaneously co-registers the structural MRI scans (T1, T1-Gd, T2, T2-FLAIR images) of each patient to the reference atlas, and segments the brain tissue, including tumorous regions of peritumoral edema, enhancing tumor and non-enhancing necrosis core. As a result of applying this method to patient scans, the images and segmentations are warped into a common atlas space, accounting for the mass effect of the tumor.

#### 4.6.2 Radiomic Feature Extraction

The masks of tumor segmentations, i.e., ET, NC, ED, and WT were overlaid on each of the MRI scans or the DTI or DSC-derived parametric maps (T1, T1-Gd, T2, FLAIR, DTI-AD, DTI-FA, DTI-RD, DTI-ADC, DSC-rCBV, DSC-PH, DSC-PSR, DSC-PC1 to PC5) to extract radiomic features of shape, volume, intensity, first-order histogram, gray-level co-occurrence matrix (GLCM), gray-level run-length matrix (GLRLM), gray-level size zone matrix (GLSZM), neighborhood gray tone difference matrix (NGTDM), local binary pattern (LBP), and Collage features. The extracted radiomic features were normalized using z-scoring. Furthermore, for each patient, *n* = 9 loci features were generated by overlaying the tumor core (ET+NC) on nine brain regions, i.e., frontal, temporal, parietal, and occipital lobes, basal ganglia, insula, corpus callosum, cerebellum, and brain stem, and calculating the average probability of the tumor core belonging to each of these regions. For *n* = 228 subjects that had all the conventional and advanced MRI scans, a total of *n* = 5,569 features, and for *n* = 358 subjects that only had conventional MRI scans, *n* = 2,477 features were extracted from multi-parametric MRI scans.

Pairwise correlation was performed on the extracted radiomic features using Pearson’s method and in highly correlated pair of features (*r* > 0.85), one of the features was eliminated. The following model training methods were carried out in the discovery cohorts for each of the classification scenarios, and the trained model was independently tested on the replication cohorts.

#### 4.6.3 Deep Learning-Based Radiogenomic Signatures

In the discovery cohort, the remaining radiomic features were ranked using recursive feature elimination with random forests and the optimal number of features was determined through nested cross-validation (10 folds for the outer and 5 folds for the inner folds). The selected feature subset was imported into the CNN classifier, as described below, for prediction of mutation status in the key driver genes, i.e., *EGFR*, *NF1*, *PTEN*, and *TP53*, and pathways, i.e., *RTK* and *PI3K*.

In the discovery cohort, we trained a CNN classifier based on a modified 34-layer ResNet architecture ^61^, with an initial 7×7 convolution and four layers (layers 1 to 4, composed of 3, 4, 6, and 3 residual blocks respectively; each residual block comprises two 3×3 convolution), as proposed by Choi *et al* ^62^. Slices of two structural images, i.e., T1-Gd and T2, along with slices of tumor segmentations (overall three image volumes) were passed as inputs for training the ResNet model. Specifically, the structural images and segmentation masks were cropped with margins of 8 pixels from the boundaries of brain tissue and resampled to 128×128×128 voxels. Intensity normalization of the brain tissue was performed using z-scoring for all structural MRI scans. Five axial slices were identified from the tumor segmentations, where the axial slice in the center included the maximum tumor mask area; 2 upper and 2 lower slices were also identified along with the center slice. After concatenating all three volumes together, the five slices were finally extracted from this concatenated volume. The discovery cohort was split into 70%-30% training-validation sets. The ResNet model was pretrained with input size of 5×3×128×128×128 for each subject in the training set (Supplemental Information, SI2, **Figure S1**). After pretraining the ResNet, the weights from the initial 7×7 convolution to layer 3 were fed to the CNN classifier and were not used in any further training process.

The radiomic features (including spatial location features) were imported as inputs to the fully connected layers that followed layer 4, and along with the weights from layer 4 were used to train the CNN classifier. Input images were further augmented using rotation. Optimization was performed using Adam optimizer (initial learning rate (*x0*) = 0.0001), with *L2* penalty of 0.01, the cross-entropy cost function, and an exponential learning rate decay (*xepoch* = *x0*\*0.99^epoch^). For both pretraining and training phases, the maximum training epoch was set to 75, and the model was saved when minimum validation loss was achieved.

#### 4.6.4 Machine Learning-Based Radiogenomic Signatures

We also trained a classical machine learning model, using support vector machines (SVM), with radiomic features. For the patients in the discovery cohort, feature selection and classifier training were performed using Least Absolute Shrinkage and Selection Operator (LASSO) feature selection approach wrapped with SVM with a linear kernel. This method was applied in a nested cross-validation (nested-CV) schema with 5-fold CV in the inner loop for feature subset selection, model optimization and hyperparameter selection, and with 10-fold CV in the outer loop to avoid data overfitting and ensure generalizability.

#### 4.6.5 Classification Scenarios

Radiogenomic signatures of co-occurring mutations in key driver genes, i.e., *EGFR*, *PTEN*, *TP53*, *NF1*, and key pathways, i.e., *RTK*, *PI3K*, *P53*, and *MAPK* were generated based on (1) radiomic features extracted from multi-parametric conventional MRI scans using deep learning and SVM classification methods, and (2) the features calculated from multi-parametric conventional and advanced MRI scans using SVM classification.

In all classification scenarios, the data was split into 75% discovery and 25% independent (unseen) replication cohorts. The performances of our predictive models in stratifying mutation status were assessed based on area under the receiver operating characteristic (ROC) curve (AUC) and balanced accuracy.

#### 4.6.6 Histograms of Imaging Features

Histograms of the most predictive features were generated for the mutant and wildtype tumors for each of classification scenarios, to gain a deeper understanding about the underlying biological processes that follow certain genetic mutations. Histograms indicate the frequency (y-axis) of a feature value (x-axis) based on the data from all patients.

