## Supplementary material for "The Radiogenomic and Spatiogenomic Landscapes of Glioblastoma, and their Relationship to Oncogenic Drivers": Supplemental Information.docx

### 1 Supplemental Information

SI1. **Data Overview**

Table S1. A summary of demographic and clinical characteristics and tumor genomics of the included IDH-wildtype patients

|  | **Cohort (conventional MRI)** | | **Cohort (conventional and advanced MRI)** | |
| --- | --- | --- | --- | --- |
| ***Demographics*** |  | |  | |
| *No. of Patients* | 358 | | 228 | |
| *Mean Age (years)* | 63.2 | | 62.4 | |
| *Age Range (years)* | 20.7 – 89.6 | | 22.0 – 89.6 | |
| *No. of Females* | 132 | | 94 | |
| ***Survival (months)*** |  | |  | |
| *Mean ± Std* | 15.8 *±* 14.1 | | 16.9 *±* 14.1 | |
| *No. of Censored Patients* | 92 | | 58 | |
| ***MGMT Methylation Status*** |  | |  | |
| *Methylated* | 95 | | 48 | |
| *Unmethylated* | 140 | | 73 | |
| *Indeterminate or Not Available* | 123 | | 107 | |
| ***Driver Genes*** |  | |  | |
|  | **Co-occurring** | **Exclusive** | **Co-occurring** | **Exclusive** |
| *EGFR_mut_/EGFR_wt_* | 83/274 | 36/43 | 62/166 | 25/24 |
| *NF1_mut_/NF1_wt_* | 62/295 | 13/43 | 40/188 | 10/24 |
| *PTEN_mut_/PTEN_wt_* | 144/213 | 42/43 | 86/142 | 21/24 |
| *TP53_mut_/ TP53_wt_* | 103/254 | 27/43 | 70/158 | 19/24 |
| ***Core signaling pathways*** |  |  |  |  |
|  | **Co-occurring** | **Exclusive** | **Co-occurring** | **Exclusive** |
| *RTK_mut_/RTK_wt_* | 107/250 | 45/43 | 81/147 | 32/24 |
| *PI3K_mut_/PI3K_wt_* | 185/172 | 68/43 | 114/114 | 37/24 |
| *MAPK_mut_/MAPK_wt_* | 67/290 | - | 43/185 | - |
| *P53_mut_/P53_wt_* | 105/252 | - | 72/156 | - |

*Numbers in parentheses indicate percentages. Std = standard deviation. AUC = Area Under the receiver operating characteristic (ROC) Curve. CI = Confidence Interval*

Table S2. The CNVs in the key driver pathways

| **Pathways** | **Alterations** | **No. of Patients with Alteration (Frequency)** |
| --- | --- | --- |
| ***RTK*** | ***EGFR* gain** | 127/177 (71.7%) |
|  | ***MET* gain** | 145/177 (81.9%) |
|  | ***PDGFRA* gain** | 20/177 (11.3%) |
|  | ***FGFR2* loss** | 110/177 (62.1%) |
| ***PI3K*** | ***PTEN* loss** | 37/177 (20.9%) |
|  | ***PIK3CA* gain** | 15/177 (8.5%) |
|  | ***PIK3R1* gain** | 22/177 (12.4%) |
| ***P53*** | ***TP53* loss** | 41/177 (23.2%) |
|  | ***MDM4* gain** | 18/177 (10.2%) |
| ***RB1*** | ***RB1* loss** | 3/177 (1.7%) |
|  | ***CDKN2A* loss** | 47/177 (26.6%) |

SI2. **Deep Learning Architecture**

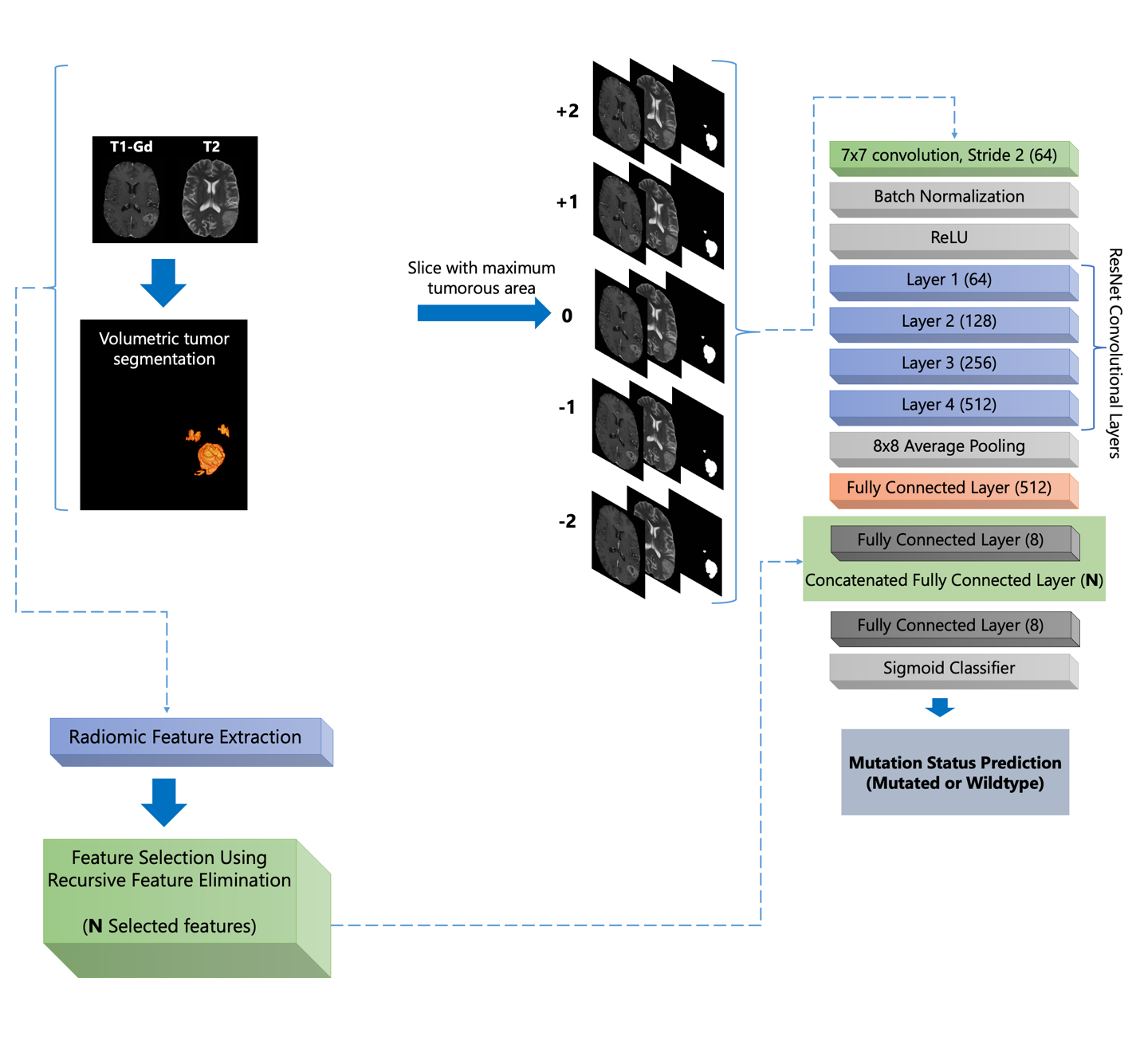

Figure S1. Deep learning model setup

SI3.
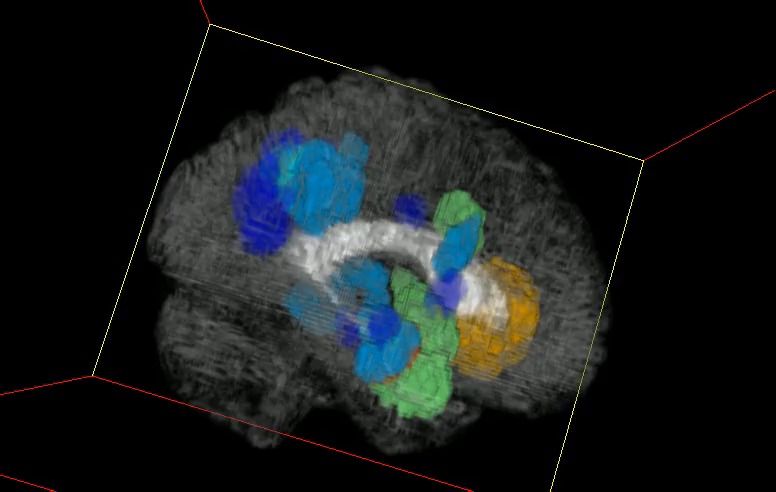
**Spatiogenomic Landscape**

The following video visualizes the 3D reconstruction of the regions of spatial location atlases (Figure 2, the right-hand side atlases in panel (A)) with most likelihood of occurrence of mutations in *EGFR, NF1, PTEN,* or *TP53* genes or their combinations. The color coding in the video is similar to those in Figure 2, panel (A).

*Supplementary Video 1. Spatial maps of frequency of occurrence for the tumors with mutations in key driver genes, including EGFR, NF1, PTEN, and TP53*

Similarly, Supplementary Video 2, provides a visualization of the 3D reconstruction of the regions of spatial location atlases (Figure 2, the right-hand side atlases in panel (C)) with most likelihood of occurrence of mutations in *RTK, MAPK, PI3K,* or *P53* pathways or their combinations. The color coding in the video is similar to those in Figure 2, panel (C).

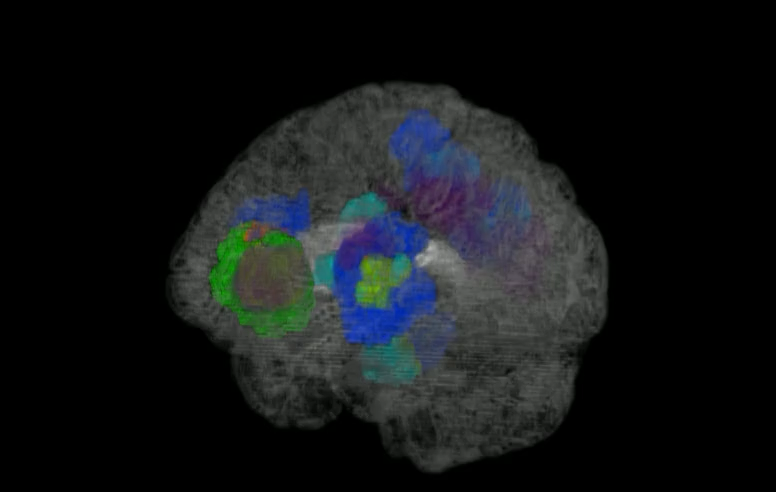

*Supplementary Video 2. Spatial maps of frequency of occurrence for the tumors with mutations in key driver pathways, including RTK, MAPK, PI3K, and P53*

SI4. **Radiogenomic Signatures**

Table S3. Performance metrics, in terms of area under the receiver operating characteristics (ROC) curve (AUC) and balanced accuracy, for radiogenomic signatures of key driver genes and pathways generated for all patients including co-occurring tumors and their values in mutually exclusive tumors. The signatures were created using Deep Learning model based on conventional MRI, and SVM classification based on only conventional or conventional + advanced MRI. The results are indicated for the discovery and replication cohorts.

|  | **Deep (conventional MRI)** | | **SVM (conventional MRI)** | | **SVM (conventional and advanced MRI)** | |
| --- | --- | --- | --- | --- | --- | --- |
|  | **AUC (95% CI)** | **Balanced Accuracy (%)** | **AUC (95% CI)** | **Balanced Accuracy (%)** | **AUC (95% CI)** | **Balanced Accuracy (%)** |
| **Discovery Cohort** | | | | | | |
| ***EGFR* (co-occurring)** | 0.79 (0.73-0.87) | 76.1 | 0.72 (0.64-0.79) | 70.3 | 0.85 (0.78-0.93) | 84.9 |
| ***PTEN* (co-occurring)** | 0.70 (0.63-0.76) | 77.8 | 0.68 (0.61-0.74) | 67.2 | 0.87 (0.81-0.93) | 82.8 |
| ***TP53* (co-occurring)** | 0.73 (0.66-0.80) | 76.5 | 0.74 (0.67-0.81) | 73.5 | 0.83 (0.76-0.91) | 81.3 |
| ***NF1* (co-occurring)** | 0.77 (0.69-0.85) | 75.3 | 0.65 (0.56-0.74) | 64.9 | 0.86 (0.77-0.95) | 83.0 |
| ***EGFR* (exclusive)** | 0.97 (0.93-1) | 98.4 | 0.83 (0.72-0.94) | 82.6 | 0.90 (0.80-1) | 92.5 |
| ***PTEN* (exclusive)** | 0.92 (0.85-0.99) | 88.9 | 0.79 (0.68-0.91) | 77.6 | 0.91 (0.79-1) | 87.1 |
| ***TP53* (exclusive)** | 0.96 (0.89-0.1) | 93.8 | 0.80 (0.66-0.94) | 85.2 | 0.77 (0.59-0.95) | 74.7 |
| ***RTK* (co-occurring)** | 0.71 (0.64-0.78) | 71.4 | 0.78 (0.70-0.84) | 76.5 | 0.84 (0.78-0.91) | 83.5 |
| ***PI3K* (co-occurring)** | 0.77 (0.71-0.83) | 75.4 | 0.69 (0.62-0.75) | 67.4 | 0.85 (0.80-0.91) | 81.9 |
| ***P53* (co-occurring)** | 0.71 (0.64-0.78) | 76.1 | 0.70 (0.63-0.78) | 69.1 | 0.82 (0.75-0.90) | 84.1 |
| ***MAPK* (co-occurring)** | 0.75 (0.67-0.83) | 69.0 | 0.72 (0.63-0.81) | 71.2 | 0.81 (0.71-0.90) | 80.8 |
| ***RTK* (exclusive)** | 0.70 (0.56-0.85) | 75.0 | 0.74 (0.63-0.86) | 75.2 | 0.82 (0.70-0.95) | 82.5 |
| ***PI3K* (exclusive)** | 0.78 (0.66-0.89) | 79.5 | 0.78 (0.68-0.88) | 80.2 | 0.84 (0.72-0.95) | 78.7 |
| **Replication Cohort** | | | | | | |
| ***EGFR* (co-occurring)** | 0.75 (0.62-0.88) | 75.0 | 0.77 (0.65-0.90) | 77.6 | 0.84 (0.71-0.98) | 87.6 |
| ***PTEN* (co-occurring)** | 0.71 (0.60-0.82) | 76.2 | 0.62 (0.50-0.74) | 63.2 | 0.86 (0.75-0.97) | 81.7 |
| ***TP53* (co-occurring)** | 0.70 (0.57-0.82) | 71.6 | 0.71 (0.60-0.84) | 72.8 | 0.79 (0.65-0.93) | 82.4 |
| ***NF1* (co-occurring)** | 0.75 (0.60-0.89) | 75.2 | 0.81 (0.68-0.94) | 75.6 | 0.85 (0.69-1) | 85.7 |
| ***EGFR* (exclusive)** | 0.93 (0.81-1) | 94.3 | 0.89 (0.74-1) | 84.3 | 0.92 (0.74-1) | 92.9 |
| ***PTEN* (exclusive)** | 0.98 (0.94-1) | 99.1 | 0.73 (0.51-0.94) | 77.3 | 0.91 (0.73-1) | 87.5 |
| ***TP53* (exclusive)** | 0.94 (0.83-1) | 94.4 | 0.91 (0.77-1) | 90.9 | 0.70 (0.40-0.99) | 75.5 |
| ***RTK* (co-occurring)** | 0.72 (0.60-0.84) | 71.3 | 0.74 (0.62-0.86) | 73.8 | 0.87 (0.77-0.98) | 88.4 |
| ***PI3K* (co-occurring)** | 0.74 (0.63-0.84) | 75.2 | 0.61 (0.49-0.72) | 64.0 | 0.78 (0.66-0.90) | 77.3 |
| ***P53* (co-occurring)** | 0.76 (064-0.81) | 78.3 | 0.75 (0.63-0.87) | 70.5 | 0.78 (0.64-0.92) | 82.4 |
| ***MAPK* (co-occurring)** | 0.82 (0.78-0.84) | 74.4 | 0.77 (0.63-0.91) | 76.8 | 0.84 (0.67-0.99) | 87.9 |
| ***RTK* (exclusive)** | 0.69 (0.15-1) | 83.3 | 0.75 (0.54-0.96) | 81.8 | 0.88 (0.69-1) | 85.4 |
| ***PI3K* (exclusive)** | 0.75 (0.28-1) | 87.5 | 0.83 (0.68-0.98) | 85.0 | 0.90 (0.73-1) | 95.0 |

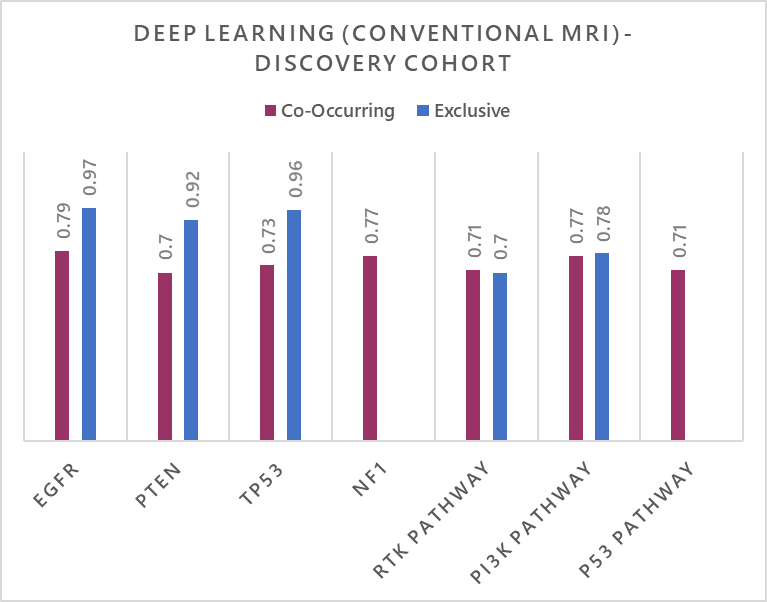

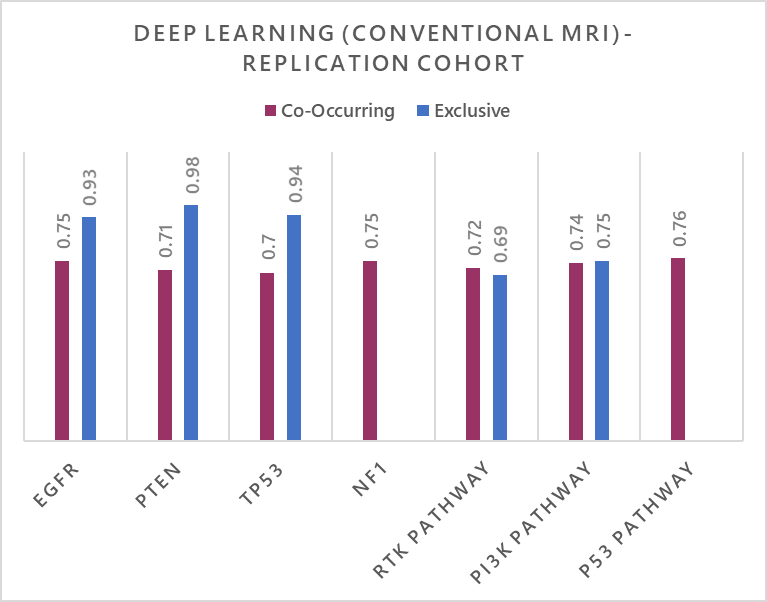

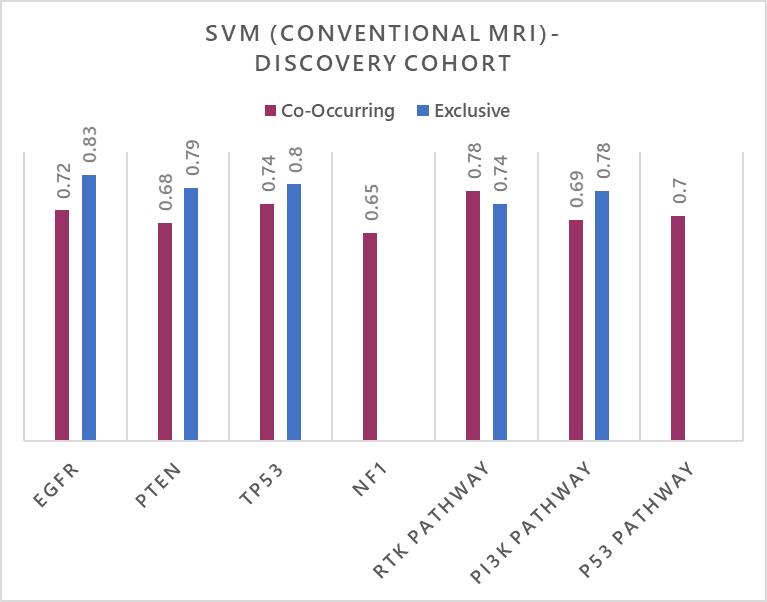

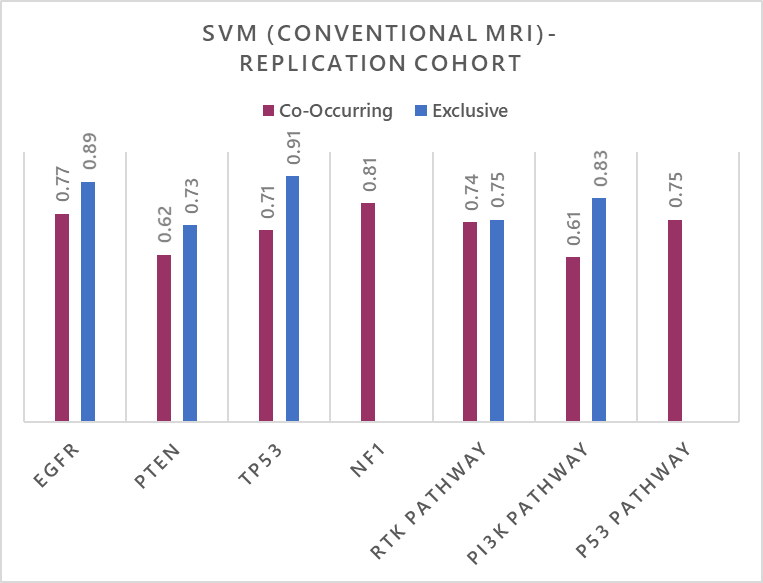

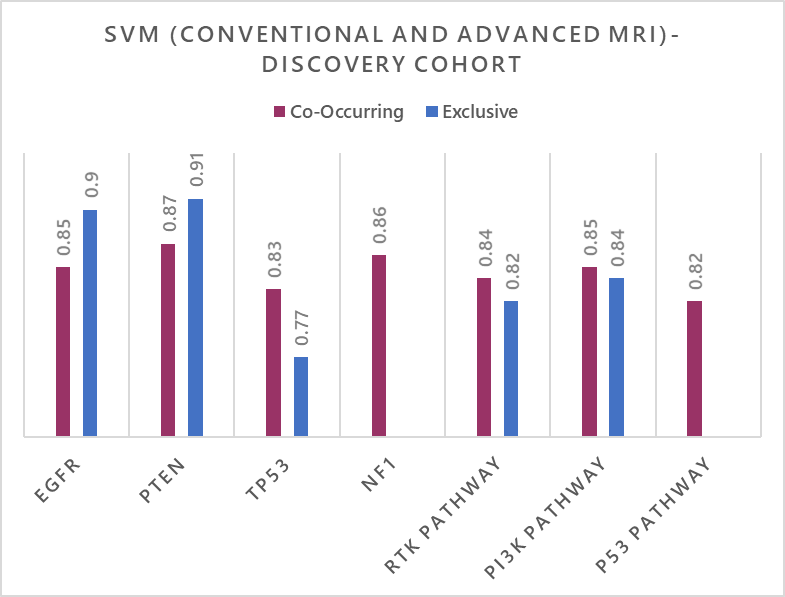

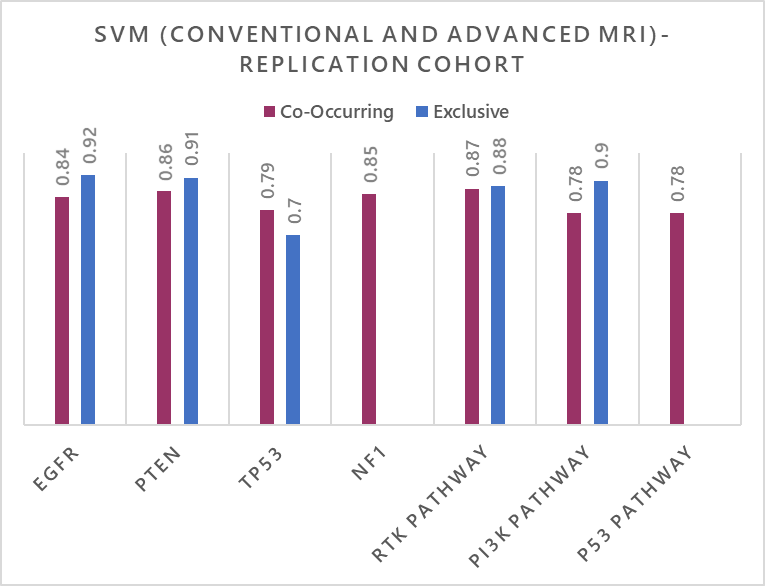

Figure S2. Bar plots displaying radiogenomic signatures generated using different machine learning/deep learning approaches as described in the manuscript. The red bars illustrate the area under the roc curve (AUC) of the co-occurring signatures and the blue bars represent the exclusive signatures.

SI5. **Evolutionary Trajectories**

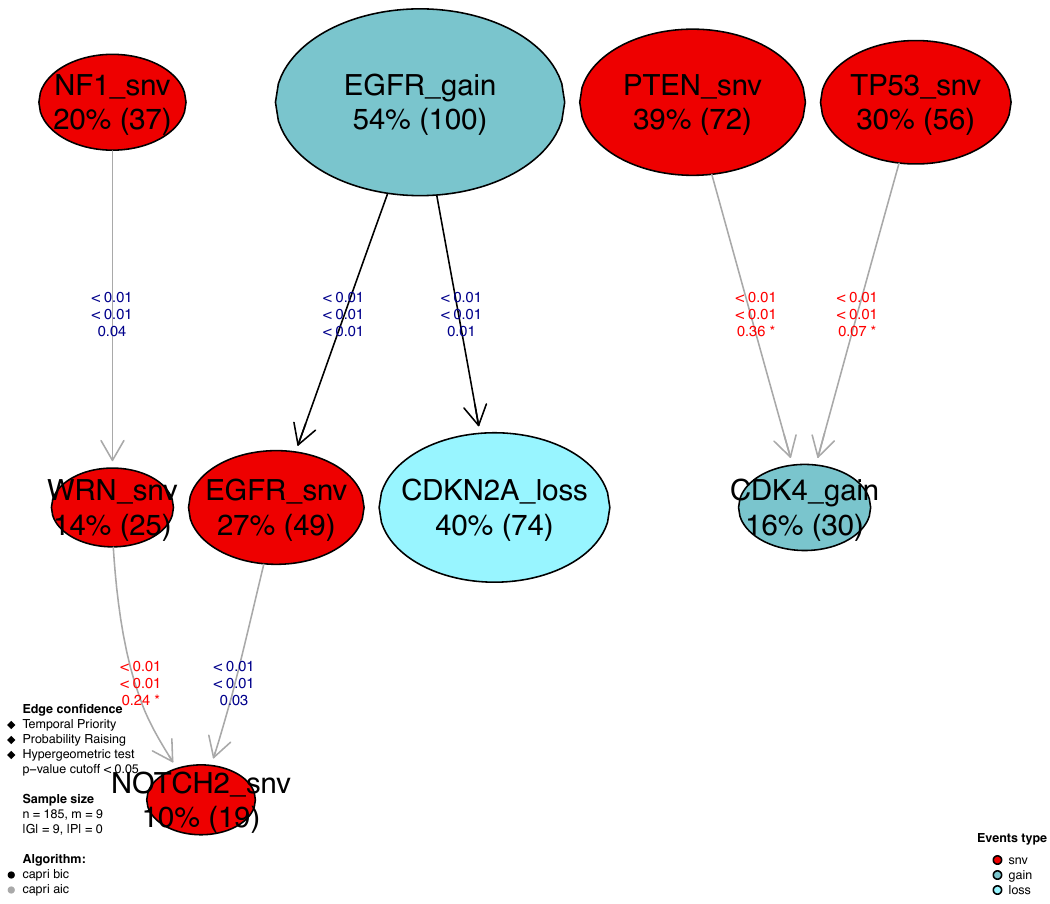

*Figure S3. Oncogenic drivers in de novo GBMs.*

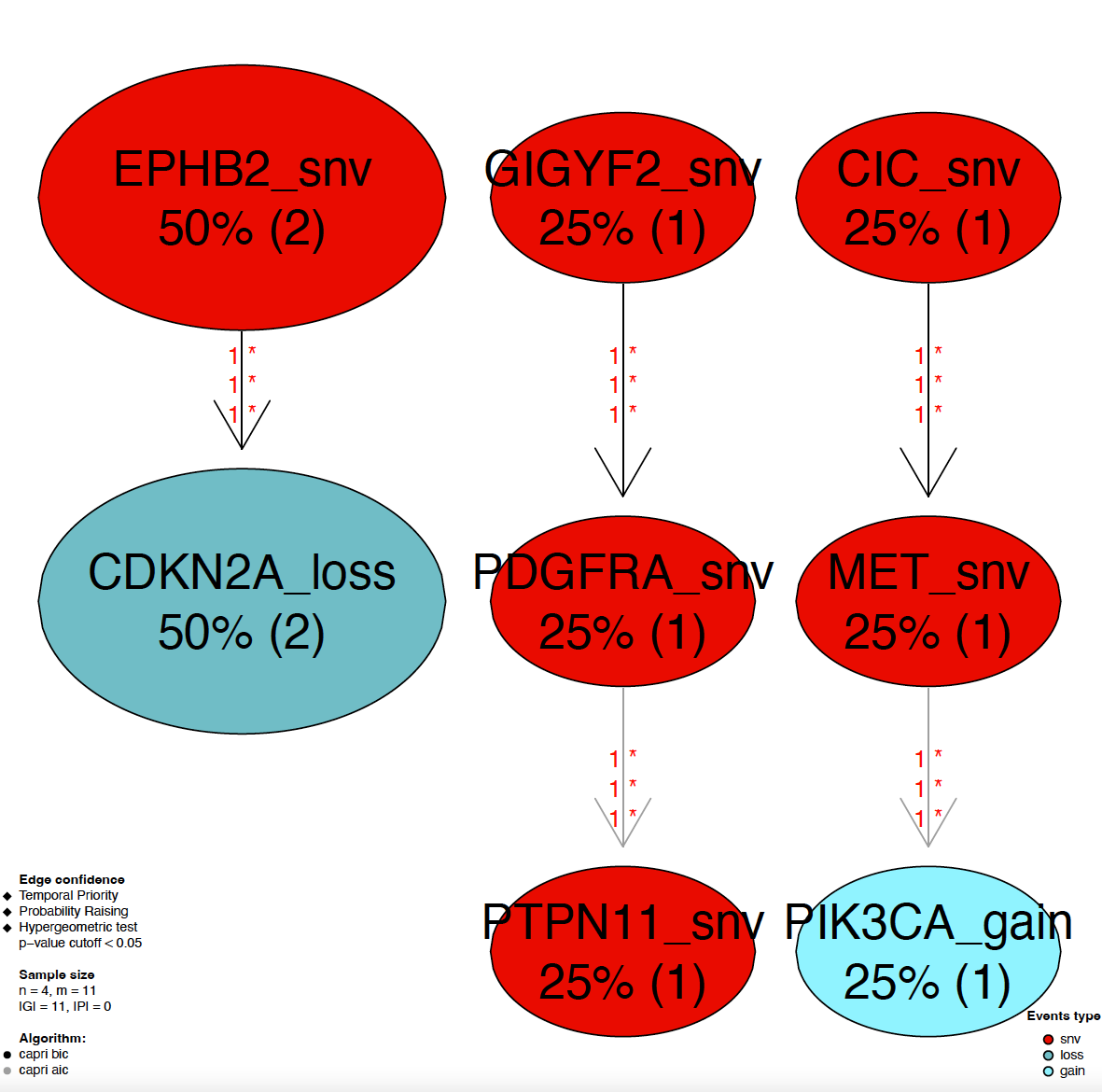

*Figure S4. Oncogenic drivers in Group 1 tumors.*

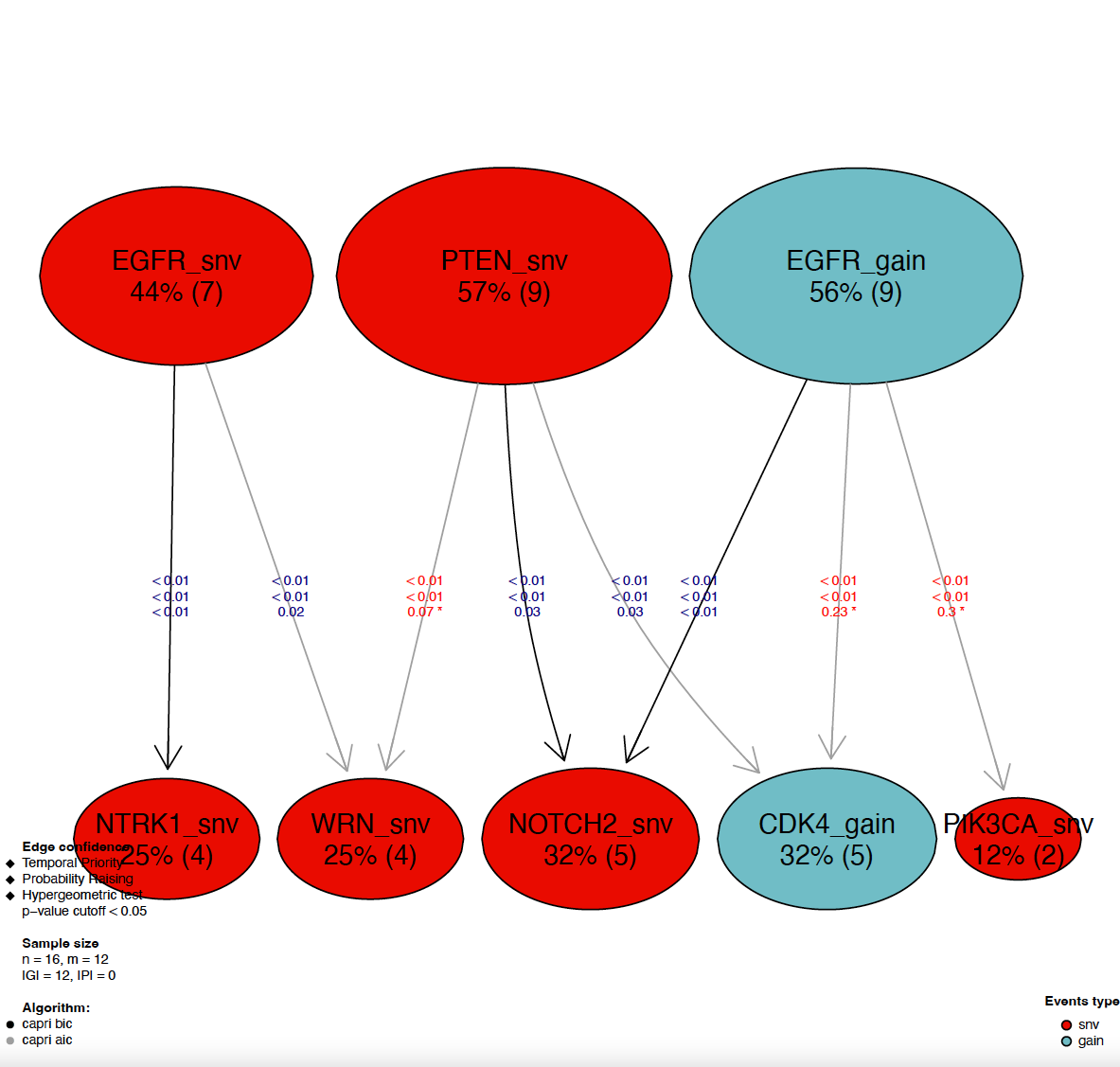

*Figure S4. Oncogenic drivers in Group 2 tumors.*

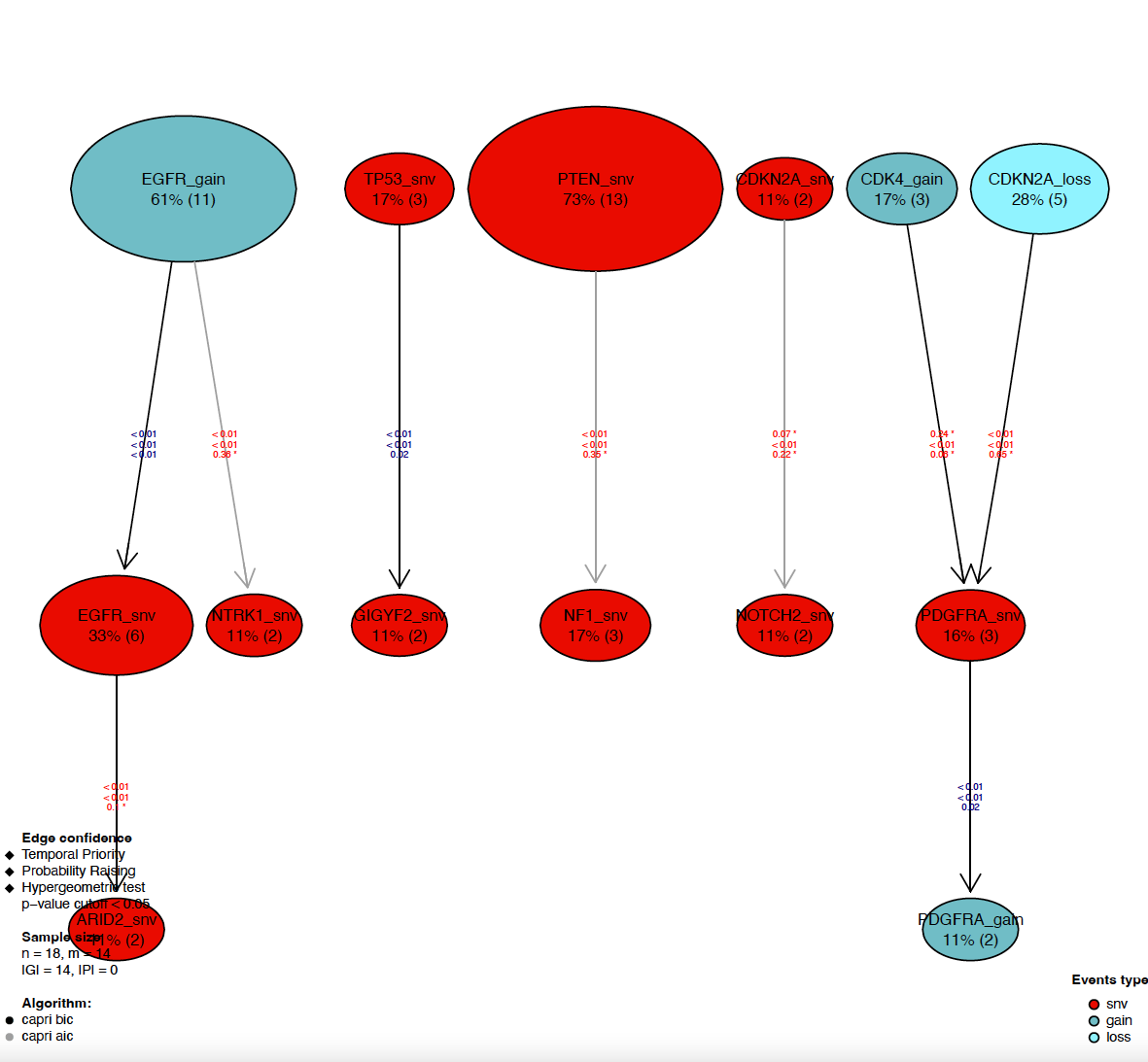

*Figure S6. Oncogenic drivers in Group 3 tumors.*

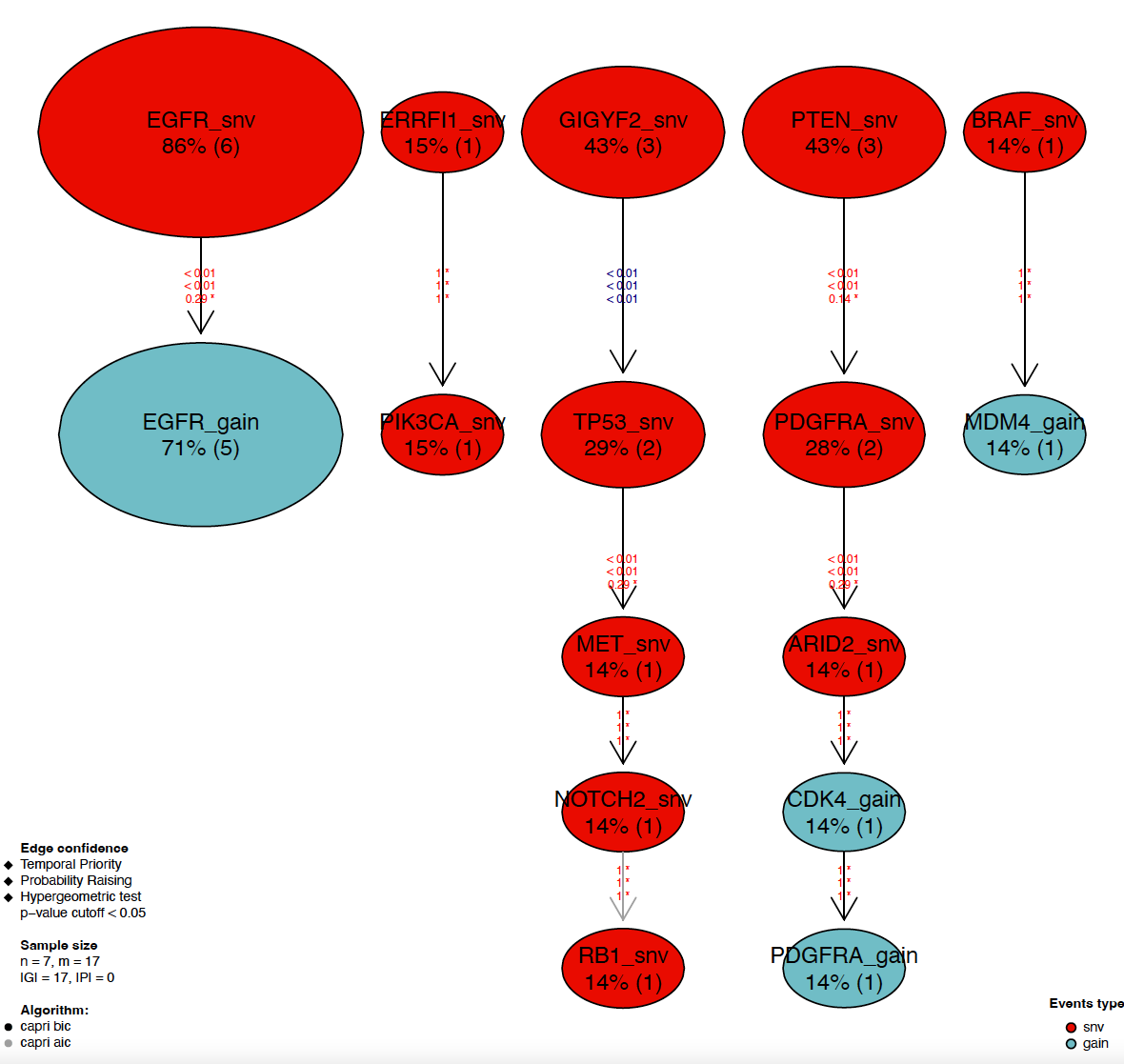

*Figure S7. Oncogenic drivers in Group 4 tumors.*
